# A newly identified detoxification system protects uropathogenic *Escherichia coli* from reactive chlorine species

**DOI:** 10.64898/2026.03.12.711259

**Authors:** Sadia Sultana, Patrick O. Tawiah, Charlie Jackson, Mehdi Bennis, Tanisha Bhimwal, Abraham Anane, Lisa R. Knoke, Alessandro Foti, Liran Paz, Mary E Crompton, Dana Reichmann, Lars I.O. Leichert, Jan-Ulrik Dahl

## Abstract

Neutrophils eliminate invading pathogens through the production of reactive oxygen and chlorine species (ROS/RCS), with hypochlorous acid (HOCl) representing the most abundant and bactericidal oxidant produced in this process. Compared to bacterial defenses against ROS, which are well studied, little is known about how pathogens respond to and counter RCS, including HOCl. Here, we identify and mechanistically characterize RcrB, a protein of the uncharacterized DUF417 protein family, for which no role in oxidative stress defense has been described yet. We report a previously unrecognized role as an RCS detoxification system that confers high-level resistance to uropathogenic *Escherichia coli* (UPEC). We show that RcrB is an inner membrane protein and strongly induced during RCS exposure and phagocytosis. Loss of RcrB results in profound HOCl hypersensitivity, accompanied by elevated macromolecular damage and severe metabolic perturbations, establishing RcrB as a central determinant of UPECs RCS stress resistance. Heterologous expression of RcrB in HOCl-sensitive intestinal *E. coli* strains is sufficient to restore resistance but requires functional glutathione biosynthesis. Quantitative HOCl trapping assays demonstrate that RcrB expression protects the bacterial population by significantly reducing extracellular HOCl, indicating active chemical quenching rather than passive membrane protection. Structure-function analysis of RcrB confirms this conclusion and demonstrate that conserved, redox-active amino acids facing the periplasm are essential for its detoxification activity. In summary, our study reveals a hitherto unknown bacterial strategy for mitigating RCS and reveal a distinct mechanism by which UPEC may survive the mammalian host defense.

**Significance Statement:** During infection, human immune cells such as neutrophils kill bacteria by releasing powerful oxidants, including hypochlorous acid (HOCl), the active ingredient of household bleach. How pathogenic bacteria survive this chemical attack is still poorly understood. This study identifies a membrane protein in uropathogenic *Escherichia coli* as a key defense factor that neutralizes HOCl before it can damage the cell. RcrB acts at the bacterial cell envelope, where it detoxifies HOCl in a glutathione-dependent manner to maintain a balanced redox homeostasis and cellular integrity, thereby even protect neighboring bacteria. These findings reveal a previously unrecognized frontline defense strategy that helps pathogens survive immune attack and may represent a new target for antimicrobial therapies.

## Introduction

The innate immune system poses one of the biggest challenges that bacterial pathogens are confronted with during the colonization of mammalian hosts. To kill bacterial intruders during infection, neutrophiles and macrophages mount the so-called respiratory burst, a host defense strategy that involves the production of highly antimicrobial reactive oxygen and chlorine species (ROS/RCS) (1–3). Superoxide generated by NADPH oxidase 2 (NOX2) in activated neutrophils dismutates to hydrogen peroxide (H_2_O_2_) (4), which is converted to hypochlorous acid (HOCl) by myeloperoxidase (MPO) (5–7). In addition, dual oxidases (DUOXs) and other related haloperoxidases generate RCS at mucosal surfaces, including the epithelial cells of the intestinal and urogenital tracts, where HOCl-dependent bacterial elimination has been demonstrated (8–10). HOCl reacts at near diffusion-limited rates with the sulfur of cysteine and methionine residues, and about three orders of magnitude slower with amino groups and metal centers (11), driving thiol oxidation (12, 13), methionine sulfoxide formation (14), *N*-chlorination of primary amines (15–17), iron–sulfur cluster disruption (18), membrane lipid oxidation (19), and ATP depletion (20). Remarkably, sub-micromolar concentrations of HOCl can eliminate vast bacterial populations within milliseconds (21). *In vivo* thiol-trapping experiments in phagocytosed *Escherichia coli* revealed pronounced cysteine oxidation and provided evidence that HOCl is the dominant antimicrobial within the complex neutrophilic ROS/RCS cocktail (22). This conclusion is also supported by genetic and pharmacological evidence: *i)* diminished NOX2 activity reduces bacterial proteome oxidation and increases susceptibility to bacterial infections in mice (22, 23); *ii)* individuals with chronic granulomatous disease exhibit heightened infection risk (5); and *iii)* MPO inhibition attenuates bacterial protein oxidation and RCS production (22). Because HOCl is short-lived and highly reactive (24, 25), its bactericidal efficacy depends on the local concentration and reaction kinetics at the bacterial cell surface, suggesting that RCS detoxification and repair pathways located on the bacterial envelope may represent important determinants for pathogen survival during host-imposed RCS stress.

Among the most common bacterial infections characterized by HOCl-mediated inflammatory responses are urinary tract infections (UTIs), which are predominantly caused by uropathogenic *Escherichia coli* (UPEC) (26). UPEC typically reside as commensals in the gut but can turn into serious pathogens upon entry to the urinary tract causing severe infections. Once UPEC ascend to the bladder, rapid infiltration of neutrophils generates a HOCl-rich milieu (27). The oxidizing environment of the bladder is likely further exacerbated by the induction of DUOX1 expression (28). UPEC also experience significant ROS/RCS exposure during pyelonephritis in the kidney (29). To successfully colonize the host, it is therefore imperative for UPEC to be able to counter RCS efficiently.

To protect themselves against RCS, *E. coli* has evolved various strategies to maintain and restore a balanced redox homeostasis and eliminate/repair the damage that these oxidants cause (18, 30–34). Among those are the activation of molecular chaperones such as Hsp33 (35), CnoX (36), RidA (37, 38), and Spy (39), which prevent the aggregation of HOCl-damaged proteins in the cytoplasm and periplasm of *E. coli*. Another strategy of *E. coli* to survive this stress is the RCS-induced conversion of ATP into polyphosphate (polyP), a protein-protective chemical chaperone (20). Bacteria also employ select transcriptional regulators, which respond to changes in RCS by increasing the expression of stress-protective target genes. Many of these transcriptional regulators can distinguish between different ROS/RCS through the specific oxidation of conserved redox-sensitive amino acid residues. Compared to hydrogen peroxide or superoxide response systems, however, which typically engage only one specific transcriptional regulator per oxidant species, at least three different HOCl-sensing transcriptional regulators have been identified in the *E. coli* K-12 strain MG1655 so far; HypT (40), RclR (41), and NemR (42, 43). Notably, what all these HOCl-sensing transcriptional regulators have in common is that their redox-regulated (in-) activation causes an elevated transcription of target genes, many of which appear to protect the organism from the toxic effects of HOCl. However, whereas bacterial defenses against H_2_O_2_ and superoxide have been extensively characterized (44–46), little is known about how these target genes protect bacteria. This is particularly relevant at the level of the cell envelope, where bacteria need to reduce the oxidant burden before widespread intracellular oxidation ensues.

We recently reported that the tolerance of UPEC to RCS is markedly enhanced compared to non-pathogenic K-12 and enteropathogenic *E. coli* strains due to the prevalence of the *rcrARB* gene cluster in this pathotype (47). Expression of this gene cluster is controlled by the RCS-specific transcriptional repressor RcrR, which is inactivated by HOCl-induced intermolecular disulfide bond formation, leading to the de-repression of *rcrARB* (47). We furthermore demonstrated that deletion of *rcrB* sensitizes UPEC to RCS stress *in vitro* without affecting growth under non-stress conditions, and that its presence enhances bacterial survival during phagocytosis in isolated human neutrophils (47), and correlates with HOCl resistance across clinical isolates (47, 48). While these results established the RcrARB system as a potentially physiologically relevant determinant of UPEC fitness within the inflamed urinary tract, it left open the crucial question by which mechanism RcrB protects UPEC against RCS mediated damage.

Here we report that RcrB functions as an HOCl detoxification system at the interface of the inner membrane and the periplasm. We demonstrate that RcrB expression alone is necessary and sufficient to protect UPEC from HOCl by reducing the amount of extracellular HOCl and limiting its influx into the cytoplasm. This, in turn, protects bacteria against the oxidation of cytoplasmic macromolecular targets, such as proteins and DNA, preserves the redox balance, prevents ATP depletion, and limits the activation of secondary stress regulons. Quantitative HOCl trapping assays demonstrate that RcrB expression significantly reduces extracellular HOCl, indicating active quenching of the oxidant rather than passive membrane protection. Structure-function analyses revealed that conserved periplasmic lysine and tryptophan residues are required for RcrB’s detoxification activity consistent with a model in which RcrB directly reacts with or facilitates reduction of RCS. The activity of RcrB depends on the presence of a functional glutathione biosynthesis machinery, providing evidence that this reducing agent is required for the RCS detoxification activity of the protein. This work expands our knowledge of the bacterial RCS defense by identifying an active envelope-level oxidant interception system and helps to explain how UPEC can maintain their fitness in HOCl-rich inflammatory environments.

## Results

### UPEC RcrB accumulates during HOCl stress *in vitro* and phagocytosis in neutrophils

We previously identified RcrB as a key determinant of UPECs resistance towards RCS, including HOCl (47, 48). As part of the *rcrARB* operon, *rcrB* is transcriptionally de-repressed upon HOCl-mediated oxidation and inactivation of the RCS-sensing transcriptional repressor RcrR (47). While the protective role of RcrB under HOCl stress has been established, the mechanism by which RbrB performs this function was unknown. To monitor abundance and localization of RcrB in UPEC strain CFT073, we generated a RcrB-superfolding green fluorescent protein (RcrB-sfGFP) fusion under the control of its native promoter. Growth curve-based studies in the presence of increasing HOCl concentrations showed that RcrB-sfGFP fully complements the HOCl-sensitive phenotype of CFT073Δ*rcrB* cells (**Fig. 1A**), confirming that the fusion protein is fully functional. Flow cytometric analysis of Δ*rcrB* cells complemented with RcrB-sfGFP and cultivated in the presence of sublethal concentrations of HOCl for 90 mins demonstrated a time-dependent increase in RcrB-sfGFP fluorescence following HOCl exposure, reaching maximal levels around 60 min (*SI Appendix, Fig. S1A,B*). Of note, expression of sfGFP under a constitutive promoter was unaffected by the treatment, excluding the possibility of stress-induced unspecific effects on the tag. Western blot analysis using an anti-GFP antibody further supported these findings and revealed an approximately two-fold increase in RcrB-sfGFP levels under HOCl stress (**Fig. 1B**; *SI Appendix, Fig. S1C*). We have previously shown that UPEC strain CFT073 is significantly more resistant to killing by phagocytosis than the non-pathogenic K-12 strain MG1655, which in large parts depends on the presence of RcrB (47). To determine whether RcrB is upregulated in the HOCl-rich environment of the neutrophil phagosome, we infected isolated primary human neutrophils with Δ*rcrB* + RcrB-sfGFP at a multiplicity of infection of 5:1 and visualized RcrB-expressing bacteria via sfGFP fluorescence by live cell imaging. Uninfected neutrophils and Δ*rcrB* + RcrB-sfGFP cells alone were included as controls. Consistent with our flow cytometry data (*SI Appendix, Fig. S1A,B*), bacteria did not show any detectable RcrB-sfGFP fluorescence when cultivated under non-stress conditions but revealed clear RcrB-sfGFP foci when engulfed by neutrophils (**Fig. 1C**). These results, which demonstrated that RcrB accumulates in response to RCS treatment both *in vitro* as well as upon phagocytosis by isolated neutrophils, support our previous observations (47) that the increased HOCl resistance of CFT073 to phagocytosis depends, at least in part, on the presence of RcrB.

**Fig 1:**
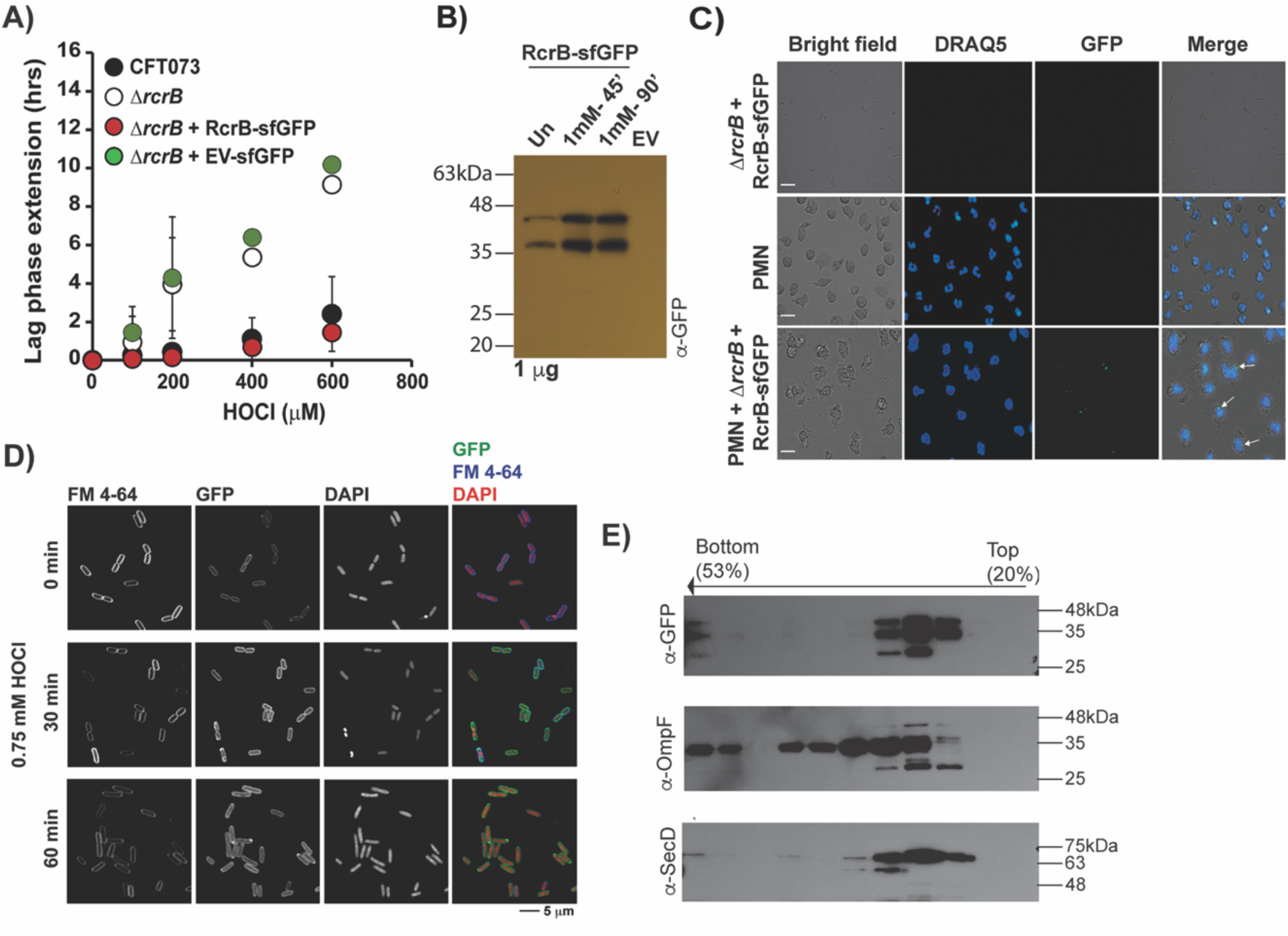
HOCl-stress induces the expression of the membrane protein RcrB. **(A)** Growth phenotype analyses of UPEC strains CFT073 (black circles), Δ*rcrB* (white circles), and Δ*rcrB* carrying RcrB-sfGFP (red circles) or the empty vector (EV) control sfGFP (green circles) were performed in MOPSg media in the presence of the indicated HOCl concentrations. HOCl-mediated lag phase extensions (LPE) were calculated for each strain (*see Materials and Methods*) (n=3, mean±S.D.). **(B)** Exponentially growing Δ*rcrB* cells carrying the RcrB-sfGFP expression plasmid or the EV control were treated with 1 mM HOCl for the indicated times. 1 µg whole cell lysate was used for western blot analysis using α-GFP antibody. RcrB: 17.1 kDa; sfGFP: 27.5 kDa. One representative image of three independent experiments is shown. **(C)** Representative epifluorescence live-cell images of isolated primary human neutrophils infected with Δ*rcrB* + RcrB-sfGFP at a multiplicity of infection (MOI) of 5:1 were imaged after 20 min of incubation. Uninfected neutrophils and bacteria alone were included as controls. Neutrophil nuclei were stained with DRAQ5 to label intracellular DNA, and bacteria were visualized via sfGFP fluorescence. Scale bar, 10 µm. Shown is one representative sample set from three biological replicates. **(D)** Subcellular localization of RcrB was determined by fluorescent microscopy. Exponentially growing Δ*rcrB* cells carrying the RcrB-sfGFP expression plasmid were treated with 0.75 mM HOCl for the indicated times. DAPI and FM4-64 are used to stain DNA and the outer membrane, respectively. The right panel shows an overlay image. One representative confocal micrograph of four biological replicates is shown. **(E)** The membrane fraction of Δ*rcrB* + RcrB-sfGFP cells was isolated by ultracentrifugation using a 20-53% sucrose gradient. RcrB-sfGFP was visualized by using a commercially available anti-GFP antibody. anti-OmpF and anti-SecD antibodies were used as markers for outer and inner membranes, respectively. One representative of three independent experiments is shown.

### RcrB localizes on the membrane

According to KEGG GENES search, the *rcrB* gene encodes a hypothetical inner membrane protein of the uncharacterized DUF417 protein family. To define its precise localization, we performed fluorescence microscopy of Δ*rcrB* cells expressing either sfGFP alone or the RcrB-sfGFP fusion. While sfGFP expressing Δ*rcrB* cells showed basal GFP fluorescence in the cytoplasm, the RcrB-sfGFP signals co-localized with the membrane marker FM4-64 under non-stress conditions (*SI Appendix, Fig. S1D*). Of note, the membrane localization of RcrB appeared to be growth phase-independent, with most pronounced protein puncta being detected at or near the cell pole in late-exponential and stationary cells (*SI Appendix, Fig. S1D*). Sixty minutes exposure of Δ*rcrB* + RcrB-sfGFP to sublethal concentrations of HOCl caused a significant increase in sfGFP fluorescence along the membrane **(Fig 1D)**, which is in agreement with our western blot analysis (**Fig 1B**). No such accumulation was observed in UPEC cells expressing sfGFP alone. To determine RcrB’s location in the cell envelope of UPEC more precisely, we fractionated the cell lysates by sucrose gradient ultracentrifugation and used antibodies against sfGFP (i.e.; anti-GFP), the inner membrane protein SecD (i.e.; anti-SecD) (49), and the outer membrane protein OmpF (i.e.; anti-OmpF) (50) for detection. RcrB was found in the same low sucrose density fractions as the inner membrane protein SecD **(Fig 1E),** indicating that RcrB is primarily located in the inner membrane. Based on these results, we concluded that UPEC cells respond to HOCl treatment with the upregulation of membrane-bound RcrB.

### Loss of RcrB amplifies the cellular response to HOCl

To define the physiological consequences of the *rcrB* deletion, we performed RNA-seq on mid-log CFT073 WT and Δ*rcrB* cells exposed to 0.75 mM HOCl. Under non-stress conditions, transcriptional differences were minimal. In contrast, compared to WT cells, we observed markedly amplified stress responses in *rcrB*-deficient cells with 733 genes at least 1.5-fold higher and 451 genes at least 1.5-fold lower expressed in the WT compared to Δ*rcrB* cells (**Fig. 2**; *SI Appendix, Table S1*). Analysis of the gene expression changes revealed that in the absence of RcrB, HOCl stress triggers a stronger upregulation of multiple functional gene classes, including antioxidant pathways such as genes encoding thioredoxin 2 (*trxC*), the glutaredoxin-like protein NrdH (*nrdH*), glutaredoxin 1 (*grxA*), and alkyl hydroperoxide reductase (*ahpC*). Oxidative damage repair systems were also more induced in the absence of *rcrB* compared to WT cells, including genes encoding molecular chaperones (*e.g.; hscC, ibpA, ibpB, dnaA*, and *spy*). Transcripts of several other stress response gene were more abundant in cells lacking RcrB, including many genes required for flagellar biosynthesis, *marRAB*, the or the diguanylase cyclase encoding gene *ydeH*. Notably, the RclR regulon (*i.e.; rclABC*), encoding a previously characterized Cu(II) reductase system (51), was among the highest upregulated transcripts in Δ*rcrB* when compared to WT. Notably, this operon contains RclC, which shows partial homology to RcrB, although we did not observe any functional complementation of Δ*rcrB* by this homolog in a previous study (48). In contrast, NADH dehydrogenase (*nuoCEFGL*), and ATP synthase genes (*atpAB*) were more abundant in HOCl-stressed WT compared to Δ*rcrB*. Likewise, taurine uptake (*tauABC*) and molybdenum cofactor biosynthesis genes (*moaBCDE, mobAB*) were more abundantly expressed in WT cells, likely because the latter are required for the activity of molybdoenzymes such as dimethyl- and methionine sulfoxide reductases that have previously been identified to have crucial protective roles during HOCl stress (14, 52). Gene enrichment analysis based on biological processes revealed that, compared to HOCl-treated WT cells, amino acid biosynthesis genes are most prominently downregulated in Δ*rcrB* cells (**Fig. 2B**), suggesting that the absence of RcrB causes increased intracellular HOCl stress and a possible defect in restoring proteostasis, which is further supported by the elevated transcript levels of antioxidant genes as well as genes involved in protein damage repair. We hypothesize that this increase in transcription is due to the increased HOCl influx in the absence of RcrB, which elevates the demand for repair systems.

**Fig 2:**
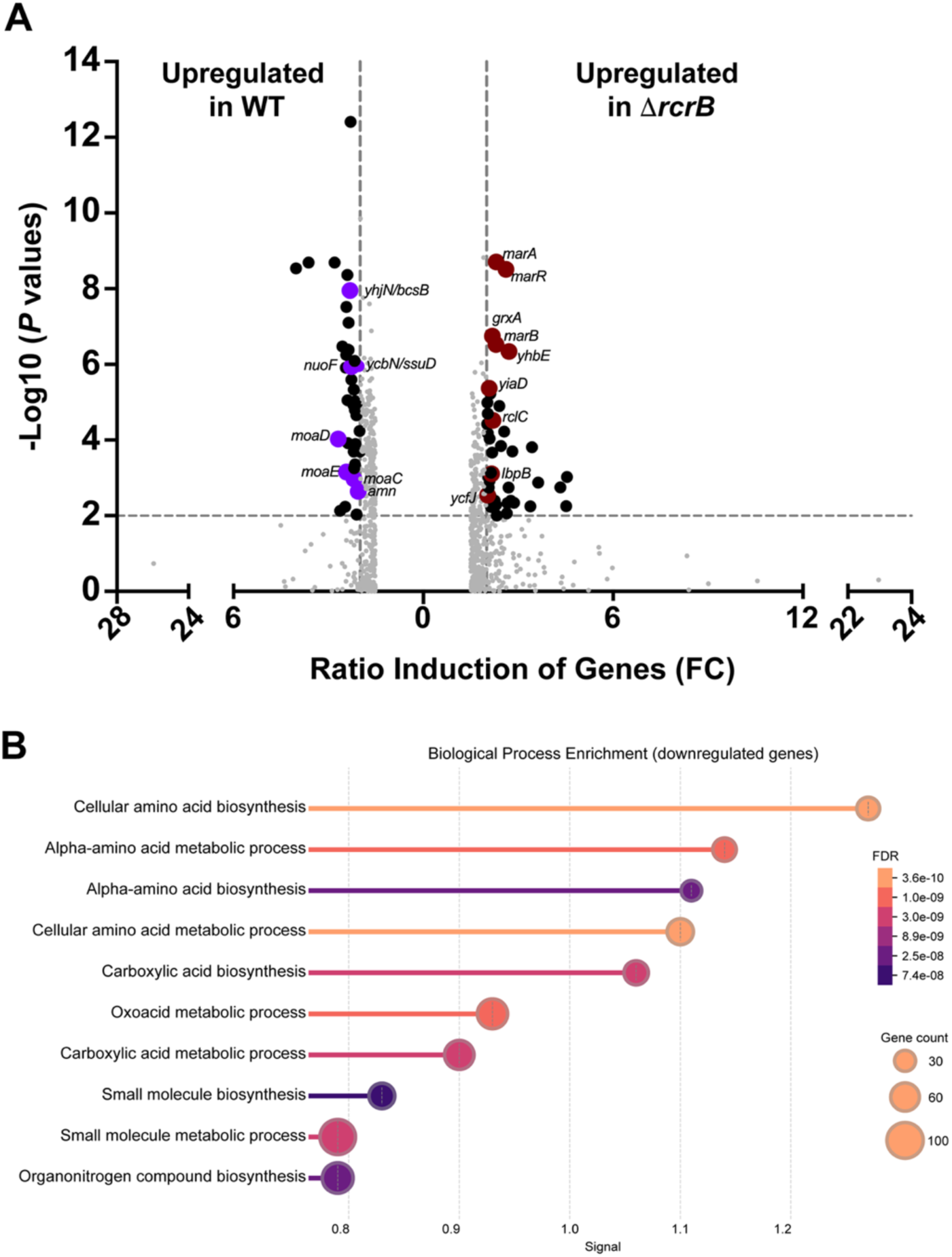
Genetic disruption of rcrB severely affects UPEC gene expression during HOCl stress. Exponentially growing CFT073 and ΔrcrB cells were incubated with and without 0.75 mM HOCl for 20 min. Transcription was stopped by the addition of ice-cold methanol before RNA was extracted and genomic DNA removed. Reads were aligned to the CFT073 reference genome (NCBI accession number: AE014075). Differences in transcript level in response to HOCl treatment was determined for CFT073 and ΔrcrB cells, respectively. **(A)** The Volcano plot illustrates the ratio of transcript levels in ΔrcrB/CFT073 (right side) or CFT073/ΔrcrB (left side). Transcript levels of genes more abundant in HOCl-treated ΔrcrB compared CFT073 are depicted as positive values, whereas transcript levels of genes more abundant in HOCl-treated CFT073 compared to ΔrcrB are depicted as negative values. Values >1.5-fold changes were plotted and >2-fold changes were highlighted (M ≥2 or ≤ −2, P≤ 0.05). Black dots: genes with fold changes >2; red dots: selected genes more induced in ΔrcrB; purple dots: selected genes more induced in CFT073. Transcriptome analysis was performed from three independent biological replicates. **(B)** Gene Set Enrichment Analysis of biological processes downregulated in HOCl-treat Δ*rcrB* compared to WT. Figure was generated using STRINGv12.0.

Quantitative real-time-PCR (qRT-PCR) analysis of stress-responsive markers for protein aggregation (*ibpA*), DNA damage (*sulA*), and lipid peroxidation (*yqhD*) confirmed that while 0.75 mM HOCl induced all three genes in both CFT073 and Δ*rcrB*, induction was significantly more pronounced in the absence of RcrB (**Fig. 3A**). Conversely, recombinant expression of RcrB in MG1655 reduced *ibpA*, *sulA*, and *yqhD* transcript levels in response to HOCl treatment (**Fig. 3B**). Moreover, chromosomal IbpA-monomeric superfolder green fluorescent protein (IbpA-msfGFP) expressed from its native promoter, which co-localizes with protein aggregates (53), revealed a clear concentration-dependent increase in fluorescence upon HOCl treatment in empty vector–containing MG1655 cells (*SI Appendix, Fig. S2A,B*), whereas expression of RcrB strongly reduced this response. these data suggest that expression of RcrB reduces HOCl-mediated protein aggregation and provide first evidence that presence of RcrB mitigates oxidative damage during HOCl stress.

**Fig 3:**
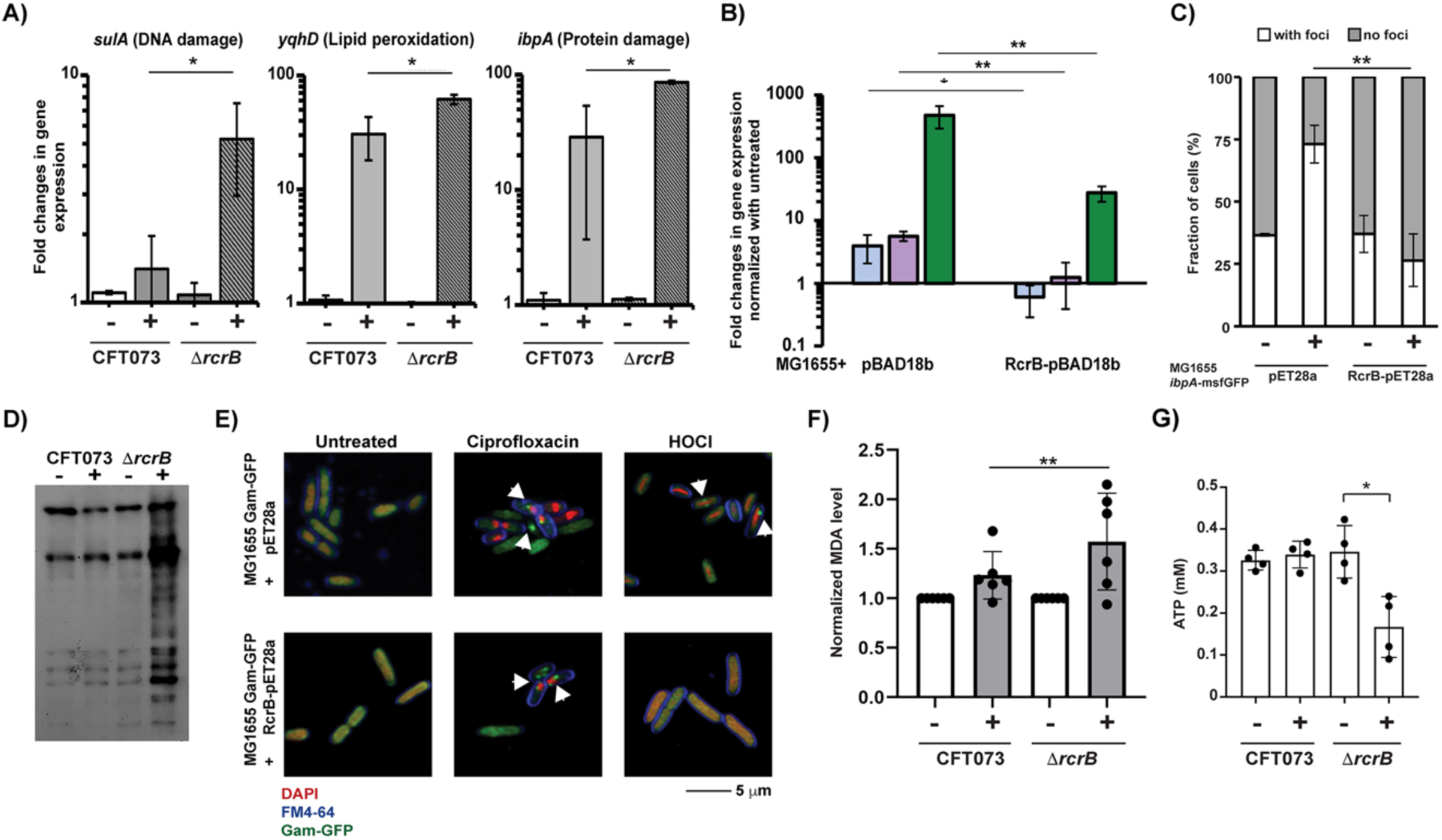
RcrB expression reduces macromolecular damage caused by HOCl. **(A,B)** The presence of RcrB alleviates the HOCl-induced expression of the stress response genes *sulA* (indicative of DNA damage), *ibpA* (indicative of protein aggregation), and *yqhD* (indicative of lipid peroxidation). The indicated strains were grown to exponential phase and treated with 0.75 mM HOCl for 20 min before transcription was stopped by the addition of methanol, RNA extracted, residual genomic DNA removed, and mRNA reverse transcribed into cDNA. Changes in transcript levels were determined by qRT-PCR. **(A)** Fold-change in gene expression was determined in strains CFT073 (WT) and Δ*rcrB*. (n= 3, mean±S.D.). **(B)** Fold-change in gene expression was determined in MG1655 cells with RcrB-pBAD18b or EV control pBAD18b. *sulA*, blue bars; *yqhD*, purple bars; *ibpA*, green bars. (n= 3, mean ± SD). **(C)** MG1655 *ibpA*-msfGFP cells transformed with either RcrB-pET28a or the EV control were grown in MOPSg to early exponential phase and incubated with HOCl for 30 min. IbpA-msfGFP foci formation was then visualized by confocal microscopy. DAPI and FM4-64 are used to stain DNA and outer membrane, respectively. Cells with RcrB-sfGFP foci were blind-counted (n=4; mean±S.D.). **(D)** Protein carbonylation was analyzed by immunoblotting in CFT073 and Δ*rcrB* cells before (−) and after (+) treatment with 0.8 mM HOCl. Shown is one representative image of three independent biological replicates. **(E)** Exponentially growing cells of the MG1655 Gam-GFP reporter strain carrying either RcrB-pET28a or the EV control were treated with and without 1 mM HOCl and 25 µg/mL ciprofloxacin (Cipro) for 2 hrs, respectively. Gam-GFP foci were visualized under the microscope and quantified by blind-counting. One representative image of five independent biological replicates is shown. **(F)** Membrane lipid peroxidation was examined in CFT073 and Δ*rcrB* cells treated with or without 1 mM HOCl using the TBARS assay. MDA (nM) was calculated using a standard curve and normalized to the untreated condition (n= 6, mean±S.D.). **(G)** ATP was quantified from exponentially growing CFT073 and Δ*rcrB* treated with and without 0.5 mM HOCl for 35 min using the Bactiter Glo Cell Viability kit (Promega). One-way ANOVA (Dunnett’s post-test) between different cell samples (GraphPad Prism) *p<0.05; **0.01>p>0.001.

### RcrB protects UPEC from HOCl-mediated macromolecular damage

Direct quantification of intracellular HOCl remains technically challenging due to the high reactivity of the oxidant with many cellular targets and the formation of secondary oxidation products with antimicrobial reactivities (15, 17). We therefore assessed the physiological consequences of RcrB deficiency by measuring HOCl-induced macromolecular damage. Following up on our observation that HOCl triggers the concentration-dependent increase in IbpA-msfGFP expression, we quantified IbpA-msfGFP foci formation in MG1655 cells with and without plasmid-encoded RcrB as an indication of protein aggregate formation in living cells. Compared to non-stress conditions, the number of cells with one or more IbpA-msfGFP foci following HOCl treatment increased two-fold in the absence of RcrB (**Fig. 3C**, *SI Appendix, Fig. S2C*), consistent with the known co-localization of IbpA with protein unfolding intermediates (53). In contrast, no stress-induced increase in IbpA-msfGFP foci was observed when RcrB was expressed suggesting that the presence of RcrB diminishes HOCl-mediated protein aggregation. We obtained similar results when we increased RcrB expression using an arabinose-inducible plasmid (*SI Appendix, Fig. S2D*). To independently validate these results, we also tested CFT073 WT and Δ*rcrB* cells for protein carbonylation, an irreversible oxidative modification of susceptible amino acid side chains (54). While basal carbonylation levels were comparable between the two strains, exposure to sublethal HOCl caused substantially higher levels of protein carbonylation in Δ*rcrB* cells than the WT (**Fig. 3D**). These results further supported the conclusion that absence of RcrB increased oxidative protein damage. To test the effects of RcrB on HOCl-mediated DNA damage (55), we followed *sulA* promoter (*P_sulA_*)-luciferase activity in HOCl-treated WT and Δ*rcrB* cells, which revealed significantly increased *P_sulA_* activity in the absence of RcrB indicating enhanced DNA damage compared to the parental strain (*SI Appendix, Fig. S2F*). To show DNA damage more directly, we utilized the MG1655 Gam-GFP double-strand break (DSB) reporter strain, which forms fluorescent foci when bound to DNA double strand breaks *in vivo* (56). In exponentially growing MG1655 Gam-GFP cells (with or without RcrB plasmid), HOCl treatment triggered robust DSB formation in cells lacking RcrB, whereas RcrB expression largely abolished Gam-GFP foci formation (**Fig. 3E**; *SI Appendix, Fig. S2E*). Of note, RcrB did not attenuate DSBs induced by the fluoroquinolone ciprofloxacin, demonstrating that the protein protects DNA specifically against HOCl-stress. HOCl readily oxidizes unsaturated fatty acids (19) resulting in the production of malondialdehyde (MDA), which in reaction with thiobarbituric acid (TBA) generates red adducts and has therefore been widely used as a biomarker for lipid peroxidation (57, 58). HOCl exposure resulted in significantly higher lipid peroxidation in Δ*rcrB* cells relative to WT (**Fig. 3F**), indicating increased oxidative damage to membrane constituents in the absence of RcrB. However, other characteristics of the membrane physiology were not impacted by RcrB (*SI Appendix, Results & Fig. S3)*. Taken together, our data revealed that loss of RcrB increases HOCl-mediated damage to proteins, DNA, and lipids, suggesting that higher levels of HOCl might reach the cytoplasm when RcrB is missing.

### RcrB preserves cellular energy homeostasis under HOCl stress

We then asked whether loss of RcrB also compromises cellular energy homeostasis during HOCl stress. HOCl is known to severely impact glucose respiration and metabolic energy by inactivating metabolic enzymes (59) and inhibiting the transport of respiratory substrates such as glucose into the cytoplasm (60–62). As a result, redox balance is perturbed during HOCl-stress, often forcing a shift away from oxidative phosphorylation and altering nucleotide pools (63). We therefore quantified ATP levels and pyridine nucleotide ratios in CFT073 WT and Δ*rcrB* cells grown in the presence or absence of HOCl. Under non-stress conditions, ATP levels were comparable between strains. In contrast, HOCl-treated Δ*rcrB* cells exhibited ∼50% lower ATP levels relative to the WT (**Fig. 3G**), indicating a severe HOCl-induced energetic collapse in the absence of RcrB. Consistent with the impaired respiratory metabolism, the NADH/NAD⁺ ratio was markedly reduced in HOCl-stressed Δ*rcrB* cells (*SI Appendix, Fig. S2G*), whereas NADPH/NADP⁺ ratios were slightly increased (*SI Appendix, Fig. S2H*), suggesting that *rcrB*-deficient cells experience more severe redox stress and exhibit a higher demand for antioxidants given that NADPH is the primary electron donor for antioxidant systems such as glutathione reductase and thioredoxin (64).

### RcrB reduces extracellular HOCl and confers community-level protection

Our data suggest that RcrB might restrict intracellular HOCl stress by reducing oxidant influx into the cytoplasm. We thus reasoned that its activity should measurably decrease extracellular RCS levels. To test this hypothesis, we indirectly quantified extracellular HOCl using the oxidation status of 3,3′,5,5′-tetramethylbenzidine (TMB). HOCl reacts with taurine to form the more stable taurine chloramine (17), which readily oxidizes TMB and generates characteristic color compound (65). Therefore, an increase in absorption of oxidized TMB correlates directly with the level of RCS present in the media. Indeed, analysis of TMB levels in the supernatants of HOCl-treated CFT073 WT and Δ*rcrB* cultures after 15 min of stress treatment revealed approximately two-fold higher RCS levels in the strains lacking RcrB (*SI Appendix, Fig. S4A*). Complementation of the Δ*rcrB* strain with plasmid-encoded RcrB significantly reduced the extracellular HOCl concentrations, resulting in an approximately 3-fold depletion of HOCl levels relative to the EV control when fully induced (**Fig. 4A**). These results indicated that cellular RcrB levels inversely correlate with residual extracellular HOCl. We confirmed the functional consequences of the RcrB-mediated HOCl depletion by cultivating naïve bacteria in sterile-filtered spent-media of these cultures. Supernatants from HOCl-treated Δ*rcrB* cultures markedly delayed growth of the HOCl-sensitive strain MG1655, consistent with elevated oxidant presence (*SI Appendix, Fig. S4B*). In contrast, supernatants from RcrB-expressing cultures caused minimal effects, suggestive of effective HOCl neutralization prior to cultivation of these strains (**Fig. 4B**; *SI Appendix, Fig. S4C*). Lastly, co-cultivation experiments yielded in similar results, demonstrating that while in contrast to the WT, Δ*rcrB* cells were highly sensitive to HOCl exposure, they survived when cultivated and HOCl-stressed in a 1:9 mixture with RcrB-expressing cells (**Fig. 4C**). No short-term or long-term fitness defects were observed for the Δ*rcrB* strain under non-stress conditions (*SI Appendix, Fig. S4D,E*). Together, these data demonstrate that RcrB expression directly lowers extracellular HOCl concentrations, thereby limiting oxidant exposure at the population level.

**Fig 4:**
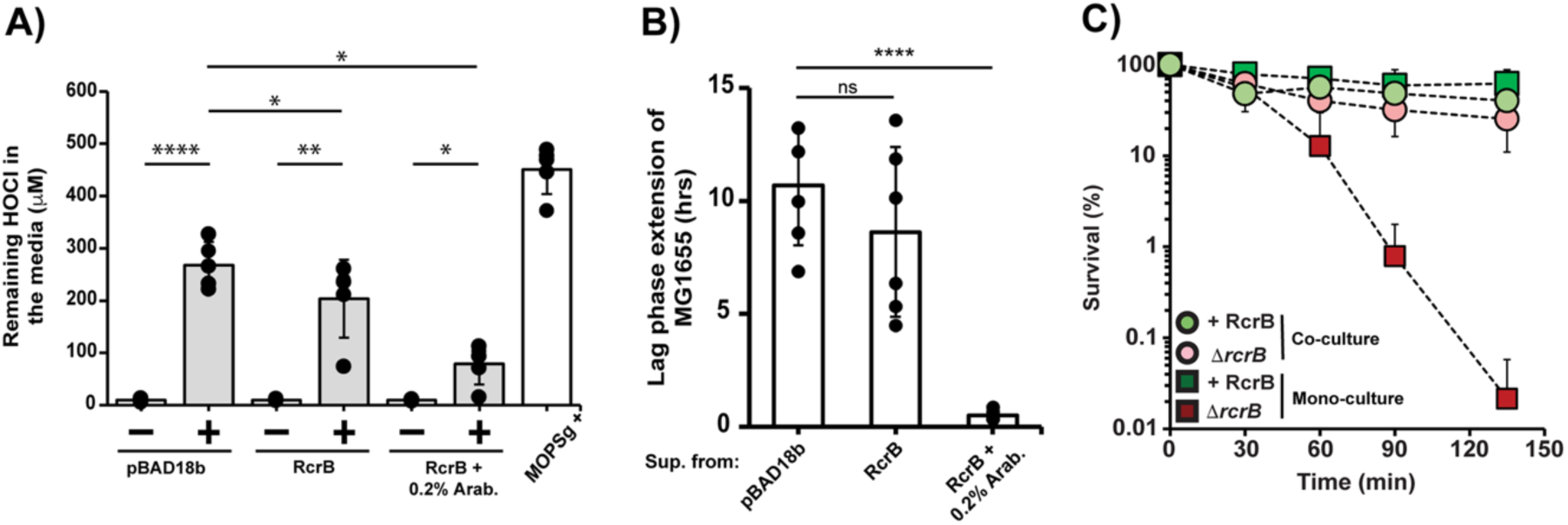
*E. coli* cellsexpressing RcrB detoxify extracellular HOCl. **(A)** Strain Δ*rcrB* containing either RcrB-pBAD18b or the EV control pBAD18b were cultivated to exponential phase in the presence and absence of 0.2% arabinose (pBAD18b, EV; RcrB, RcrB-pBAD18b without arabinose; RcrB+0.2% Arab., RcrB-pBAD18b with 0.2% arabinose). Following treatment with and without 1 mM HOCl for 15 min, cells were removed by centrifugation. Remaining HOCl and chloramines in the supernatant of the indicated strains was determined using the 3,3’,5,5’-tetramethylbenzidine (TMB) assay. (n=5, mean ± SD). Paired student t-test to compare untreated and treated conditions of each strain; and, one-way ANOVA (Dunnett’s post-test) between different treatments (GraphPad Prism) *p< 0.05, ** p<0.01, ***p<0.001, ****p<0.0001. **(B)** Growth phenotype analyses of MG1655 cells cultivated in sterile-filtered spent media from **(A)**. LPE was determined as described in *Materials and Methods* (n=6 ± S.D.). One-way ANOVA (Dunnett’s post-test) between different cell samples (GraphPad Prism) ****p<0.0001. (**C)** Overnight cultures of Δ*rcrB* (red symbols) and Δ*rcrB* containing RcrB-pBAD18b (+RcrB, green symbols) were grown overnight in the presence of 0.2% arabinose, sub-cultured into fresh MOPSg+0.2% arabinose and grown until OD_600_= 0.1. Individual monocultures (squares) were treated with 1.25 mM HOCl for 135 min and survival was examined at the indicated time points. Likewise, survival of each individual strain was assessed during co-incubation (circles) at a 1:9 ratio (Δ*rcrB:*+RcrB) following treatment with 1.25 mM HOCl. Samples were serially diluted at the indicated time points and plated onto LB agar with and without ampicillin to discriminate between cells with and without RcrB. Percent survival was calculated relative to the untreated controls (n=3-5, mean±S.D.).

### Conserved periplasmic residues are essential for the HOCl-detoxifying activity of RcrB

The data presented so far support a model in which RcrB detoxifies extracellular HOCl and limits HOCl influx into the cell. To identify the structural determinants required for this activity, we used AlphaFold prediction analysis, which posited that RcrB contains four transmembrane helices with both N- and C-termini facing the cytoplasm (**Fig. 5A**, *SI Appendix, Fig. S5*). The monomeric and dimeric states received the highest confidence scores, and the top models for each state were highly similar (*SI Appendix, Fig. S5*). We noted that several of the highly conserved residues that directly face the periplasm or are located at the interface of membrane and periplasm have redox-reactive side chains, including Lys43, Trp44, Trp60, and Lys128 (**Fig. 5B**). To test whether these residues contribute to the observed HOCl detoxification, we generated RcrB-sfGFP variants with substitution mutations in each of these amino acid residues and assessed their ability to complement the HOCl sensitivity of Δ*rcrB* cells. Indeed, substitution of Lys43, Trp44, or Lys128 completely abolished the ability of RcrB to restore HOCl resistance (**Fig. 6A**), whereas Trp60Ser exhibited only a partial defect (**Fig. 6B**). Importantly, both expression and localization of Lys43Ala, Trp44Ser, Trp60Ser, and Lys128Ala variants were comparable to WT RcrB (**Fig. 6D**; *SI Appendix, Fig. S6B*). Of note, substitution of other residues that have redox reactive side chains, are located at this interface but are not highly conserved, such as Met50 and Lys121 yielded in proteins that fully restored the HOCl resistance of Δ*rcrB*, indicating that these are irrelevant for the HOCl detoxification activity of RcrB (*SI Appendix, Fig. S6A*). Extracellular HOCl depletion assays confirmed these results and showed that supernatants from Δ*rcrB* cells expressing Lys43, Trp44 or Lys128 variants were unable to detoxify HOCl and retained HOCl levels comparable to vector controls (**Fig. 6C**). Together, our data identified conserved periplasm-facing residues, particularly Lys43, Trp44, Trp60, or Lys128 as essential determinants of RcrB function.

**Fig 5:**
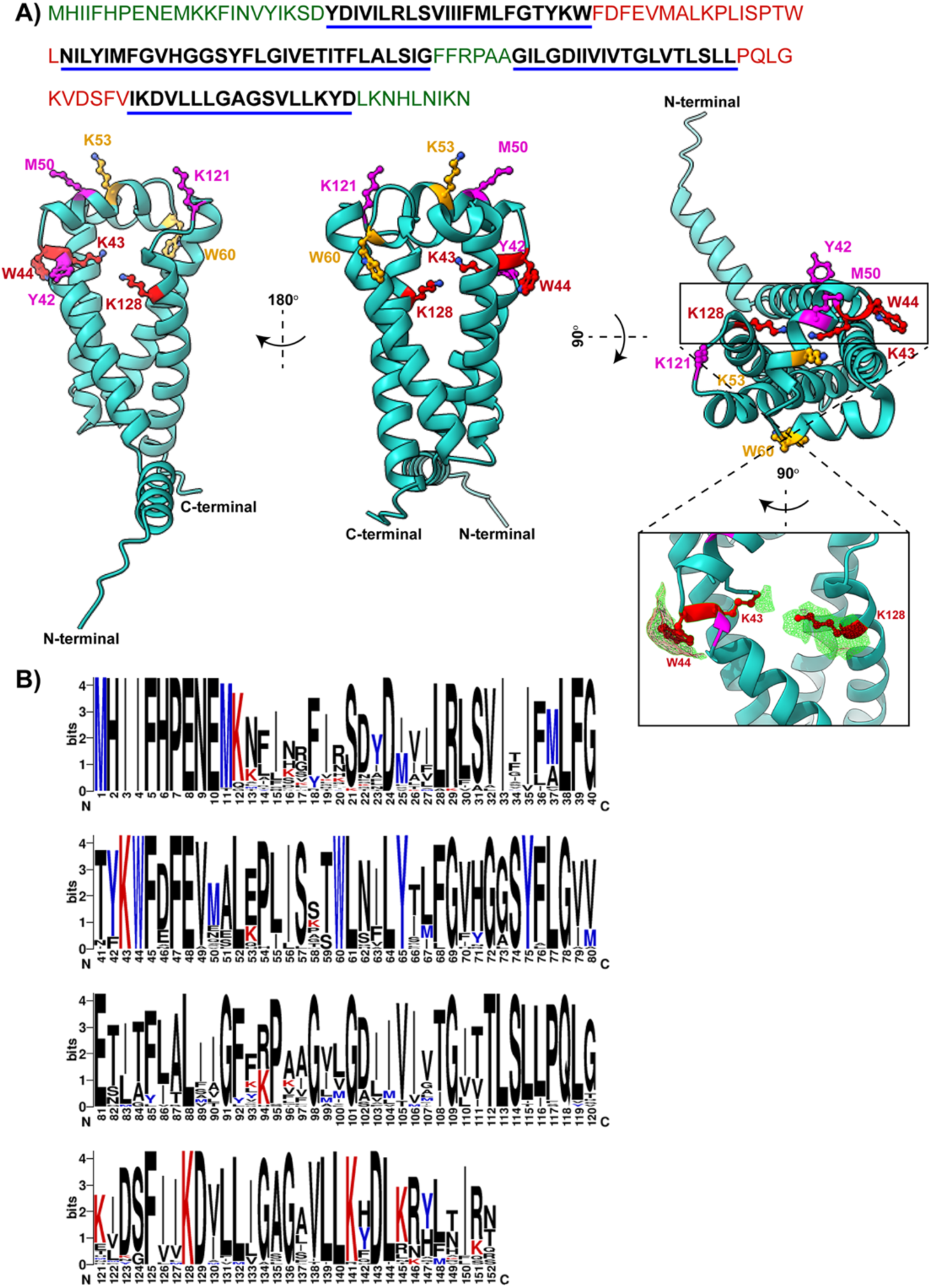
RcrB membrane topology and predicted amino acids localization. **(A)** Top: amino acid sequence of RcrB. Green: cytoplasmic; black underlined in blue: transmembrane region; red: periplasmic. Bottom: prediction by AlphaFold and Phyr2-based homology modeling. RcrB consists of four transmembrane regions. The region facing the periplasm contains several redox-active amino acids, which are highlighted in different colors depending on their significance for RcrB function (Fig. 6A**,B**; **Supplementary Fig. S6A**): yellow (intermediate impact), magenta (no impact) and red (high impact). Cartoons were created using ChimeraX. **(B)** Weblogo plot of RcrB homologs from 76 bacterial strains revealed conserved lysine (K), tyrosine (Y), and tryptophan (W) residues.

**Fig 6:**
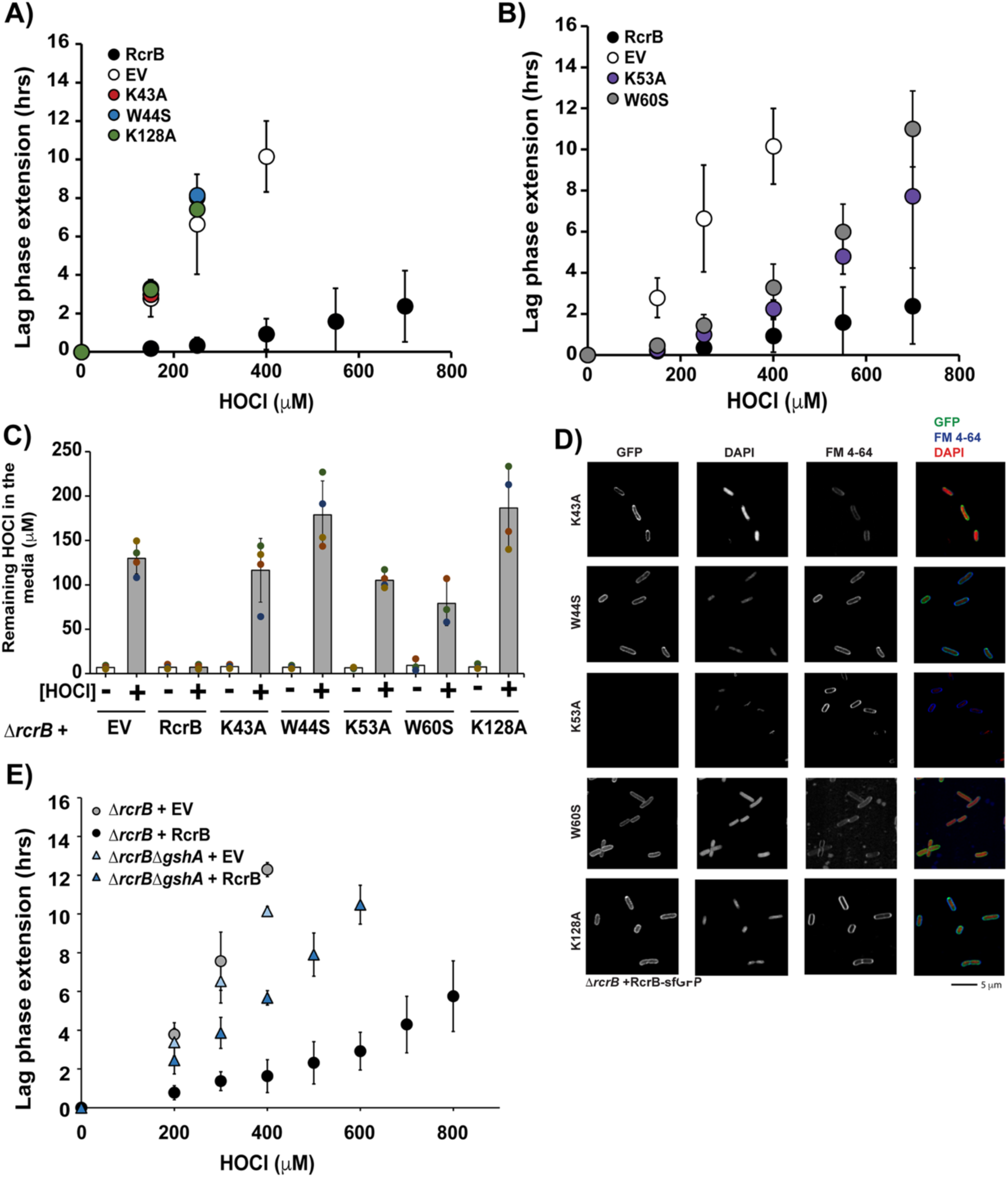
Periplasmic lysine and tryptophan amino acids are important for RcrB’s function. **(A,B)** Complementation analyses of the Δ*rcrB* strain expressing RcrB-sfGFP variants with the indicated amino acid substitutions were performed in MOPSg medium in the presence of the indicated HOCl concentrations. EV= empty vector. HOCl-mediated LPE was calculated for each strain (*see Materials and Methods*) (n = ≥3, mean± S.D.). **(C)** Plasmid-complemented Δ*rcrB* strains expressing RcrB-sfGFP variants were cultivated in the presence of 1.25 mM HOCl for 15 min, and the remaining HOCl in the medium was quantified (n=3-5, mean±S.D.). **(D)** Representative confocal micrograph of four biological replicates shows the localization of the indicated RcrB-sfGFP variant in the membrane of exponentially growing Δ*rcrB* cells. DAPI and FM4-64 were used to stain DNA and membrane, respectively. The right panel shows an overlay image of GFP, FM 4-64, and DAPI fluorescence. **(E)** Growth analyses of UPEC strains *ΔrcrB* and *ΔrcrBΔgshA* with and without RcrB from the pBAD18 expression plasmid were conducted in MOPSg medium in the presence of the indicated HOCl concentrations. HOCl-mediated LPE was calculated for each strain (*see Materials and Methods*) (n = ≥3, mean± S.D.).

### Glutathione biosynthesis Is required for RcrB function

Our data suggest a direct involvement of RcrB in HOCl detoxification at the interface of plasma membrane and periplasm. We now wondered whether the activity of the protein depends on cellular antioxidant systems. We reasoned that if RcrB functioned through reversible oxidation/chlorination reactions, its redox state would need to be continuously regenerated and hence require a potent reducing system. In Gram-negative bacteria, the most abundant thiol-based antioxidant is glutathione (GSH), which cells can readily secrete and import to balance redox homeostasis (66). We therefore constructed an Δ*rcrB* strain lacking γ-glutamylcysteine ligase (*gshA*), the enzyme catalyzing the first step of the GSH biosynthesis and tested its HOCl sensitivity in the absence and presence of plasmid-encoded RcrB. We found that the deletion of *gshA* abolished the ability of plasmid-encoded RcrB to restore HOCl resistance in Δ*rcrB* cells, which we also confirmed in the non-pathogenic K-12 strain (**Fig. 6E**; *SI Appendix, Fig. S7*). Importantly, loss of *gshA* alone did not substantially alter the growth under non stress conditions, indicating that the GSH deficiency specifically impaired RcrB-dependent protection. Deletion of *gor* (encodes the NADPH-dependent GSH reductase Gor) had no effect on RcrB activity (*SI Appendix, Fig. S7*). These findings demonstrate that GSH is required for RcrB function, likely due to the ability of the tripeptide to reduce oxidative modifications of RcrB during HOCl-stress. RcrB activity is hence not consistent with a purely sacrificial mechanism but rather an active detoxification mechanism that requires GSH to sustain RCS detoxification at the bacterial cell envelope.

## Discussion

Our study identifies RcrB as an efficient membrane-located cellular detoxification system of HOCl that reduces the influx of RCS into the cytoplasm. As a result, cytoplasmic proteins, the main targets of HOCl (64), remain protected from oxidation, providing a mechanistic explanation for the role of RcrB as the main driver of UPECs superior RCS resistance (47). Our findings converge on four central conclusions: *(i)* RcrB, which is induced during HOCl stress and phagocytosis in isolated human neutrophils on both transcriptional (47) and translational level, localizes to the inner membrane; *(ii)* lack of RcrB during HOCl stress severely impairs cytoplasmic macromolecules causing a profound imbalance in redox homeostasis and metabolism; *(iii)* RcrB actively detoxifies HOCl, thereby conferring population-level protection; and *(iv)* structure-function analyses reveal a redox-active, likely dimeric membrane protein whose activity depends on specific periplasm-facing residues and functional glutathione biosynthesis. Many RCS response and defense systems have been identified in *E. coli*, including several molecular chaperones that prevent HOCl-induced aggregation of misfolded proteins in the cytoplasm (20, 35–38) and periplasm (39), damage repair systems produced by the (in-)activation of HOCl-sensing transcriptional regulators (40–43) and two-component systems (67), and HOCl-sensitive proteins such as YdeH that switch the lifestyle of *E. coli* from planktonic to biofilms (68). Given that almost all of them are localized in the cytoplasm, they can only act after the highly reactive RCS have passed through the plasma membrane. In contrast, RcrB, which is also transcriptionally induced (47, 48), is already present at basal levels and acts directly at the periplasm-plasma membrane interface to effectively prevent intracellular oxidative damage before it occurs.

### Induced by HOCl during phagocytosis, RcrB is stably embedded in the inner membrane

We demonstrate that, while RcrB is present at basal levels, protein abundance rapidly increases upon HOCl exposure and remains stable over time, which is in agreement with our previously reported transcriptional data of RCS-specific *rcrARB* induction (48). In contrast, transcription of other HOCl defense systems such as the NemR and RclR regulons was reported to decline much earlier (41, 42, 48). Fluorescence microscopy and sucrose gradient fractionation revealed homogeneous localization of RcrB to the inner membrane as well as confirmed enhanced protein levels during HOCl exposure that remain stable over time. These data suggest that RcrB does not redistribute upon oxidative stress but instead functions at the interface between the inner membrane and the periplasm, where the HOCl molecules enter the cytoplasm. Given that HOCl is membrane-permeable and thus rapidly penetrates the cytoplasm, the membrane localization of RcrB is ideally suited for intercepting the oxidant before it can cause intracellular damage. Importantly, membrane integrity itself is not significantly altered by RcrB, consistent with reports that HOCl toxicity primarily manifests through intracellular thiol and protein oxidation rather than causing morphological disruption (64).

Expression of RcrB was also observed upon phagocytosis in isolated human neutrophils, supporting the RcrB-dependent resistance of UPEC towards neutrophil-mediated killing (47). The HOCl-generating enzyme MPO is the most abundantly expressed proinflammatory enzyme in neutrophils (1, 69), and previous reports have estimated the enzyme to be present at concentrations between 1-2 mM during inflammation in infiltrating phagocytes (5, 21, 70). A computational model predicted that MPO efficiently scavenges diffusible H_2_O_2_ at the surface of neutrophil-engulfed *Salmonella* and converts it to HOCl, which rapidly damages bacterial macromolecules (24). While HOCl is orders of magnitude more antimicrobial due to its higher rate constants with cellular thiols and methionine residues at near diffusion-limited rates, the oxidant has a ∼100-times lower lifetime compared to ROS, suggesting that its reaction with target molecules is more locally confined (64). It is therefore not surprising that bacteria have evolved a specialized system that counters HOCl directly at the cell envelope before cytoplasmic components are harmed. It is also possible that UPEC encounters HOCl outside the neutrophil phagosome, as 30% of the total cellular MPO are secreted or released into the extracellular environment (71), where DUOX-generated H_2_O_2_ may be sufficiently available to drive substantial extracellular HOCl production (9).

### Loss of RcrB amplifies macromolecular damage and disrupts redox homeostasis

We demonstrate that RcrB expression and HOCl resistance are tightly linked: deletion of *rcrB* severely compromises survival under HOCl stress, whereas chromosomal or plasmid-borne expression restores resistance. Our RNA-seq, biochemical assays, and imaging analyses collectively show that the absence of RcrB correlates with greater oxidative injury likely due to increased RCS influx into the cytoplasm. In line, *rcrB*-deficient cells accumulate higher levels of DNA damage, show a higher extent of protein carbonylation and aggregation, and exhibit a pronounced metabolic collapse, including ATP depletion and altered NAD(P)H pools. These findings align with the well-established chemistry of HOCl, which attacks purines in DNA to form mutagenic 7,8-dihydro-8-oxoguanine adducts and chlorinated nucleosides (72), resulting in slow but very efficient denaturing of DNA (63). In a double whammy, cells simultaneously depend on a functional methionine sulfoxide reductase repair system to restore the activity of the ubiquitous recombinase RecA, a cellular target of HOCl that is required for repair of damaged DNA (73). Proteins are in fact considered the primary targets of HOCl, as the side chains of several amino acids are susceptible to HOCl-mediated oxidation and/or chlorination, often resulting in widespread protein misfolding and aggregation (30, 31, 33, 34, 64, 74, 75). Consequently, *E. coli* compensates severe oxidative stress through the transcriptional upregulation of molecular chaperones, such as IbpA (42, 47), which we also have observed in this study. Other known target proteins characterized by HOCl-induced inactivation include the membrane-localized F1-ATPase (59), which was highly upregulated in HOCl-treated UPEC WT cells and could therefore explain the significant reduction in ATP levels observed in the HOCl-stressed mutant. This effect may even be exacerbated by the conversion of ATP into the chemical chaperone polyP, which has been shown to accumulate in HOCl-stressed bacteria in parts due to the oxidative inactivation of the degrading enzyme exopolyphosphatase (20, 76, 77). HOCl can also damage DNA-binding plasma membrane proteins, which leads to inhibition of DNA replication (78). Lastly, we detect RcrB-dependent imbalances in NAD(P)H ratios indicating a perturbation of NAD(P)H homeostasis in the mutant, which provides the first evidence for a link between RcrB activity and antioxidant systems, several of which were significantly more induced in HOCl-stressed Δ*rcrB* cells. Because antioxidant systems require NAD(P)H for their catalytic cycles, maintaining a cellular redox balance is essential for sustaining metabolic function and redox-buffering capacities. Thus, RcrB appears to act upstream of global oxidative collapse and the disproportionate intracellular damage observed in Δ*rcrB* cells suggests that RcrB limits effective HOCl influx by quenching the oxidant before it can reach cytoplasmic targets.

### The redox-active, likely dimeric membrane protein RcrB detoxifies HOCl and confers community-level protection

Biochemical and growth-based assays indicate that RcrB detoxifies HOCl through a redox-mediated mechanism. RcrB-expressing cells effectively eliminate extracellular HOCl, consistent with a membrane-associated detoxification mechanism that quenches HOCl before widespread intracellular damage occurs. Notably, this activity provides community-level protection, suggesting that RcrB lowers extracellular and periplasmic HOCl concentrations sufficiently to benefit neighboring cells. This might be advantageous in dense bacterial populations exposed to host-derived RCS stress during infection. Mechanistically, RcrB activity depends on GSH biosynthesis, as RcrB-mediated protection is significantly reduced in GSH-deficient backgrounds. Whether other reducing systems, such as thioredoxin, can partially complement the loss of GshA will be tested in future studies, as high redundancy has been reported for these systems (79). Our structure–function analysis identified several conserved, redox-sensitive amino acid residues on the periplasm-facing side that are essential for the activity of RcrB during HOCl stress. Substitution of Lys43, Trp44, Trp60, and Lys128 markedly compromised the ability of the protein to restore the HOCl resistance of a Δ*rcrB* strain despite preserved membrane localization and WT-like expression, demonstrating that these residues contribute directly to the detoxification mechanism rather than to protein stability. The monomeric and dimeric states received the highest confidence scores in our AlphaFold predictions, and the top models for each state were highly similar. The predicted dimer interface is formed by hydrophobic residues within the transmembrane helix, with Lys128 and Phe125 stabilizing the periplasm-facing region of the interface. If indeed RcrB is a dimer, chlorination of Lys128 could influence dimerization, either directly or by perturbing interactions with Phe125. Both Trp and Lys are well-suited to participate in redox chemistry at the membrane interface. HOCl can directly chlorinate the indole ring of Trp to form 2-oxoindole derivatives (16), or cause the generation oxygenated Trp products depending on the local amino acid sequence context (80). Much like Arg, Lys residues can be chlorinated reversibly by HOCl, which occurs in *E. coli* during the HOCl-mediated activation of the molecular chaperones RidA and CnoX (36–38, 81). Lys has also recently been proposed to have antioxidant properties, as the formation of one Lys-nitrile quenches two HOCl molecules without affecting the secondary structure of the proteins tested (82). The absence of these Lys residues, however, caused an increase in chlorinated Tyr residues, an irreversible protein modification triggering protein aggregation. The positioning and conservation of these important residues suggest that RcrB employs a coordinated redox-active surface to engage incoming HOCl at the periplasmic interface. Although RcrB contains several methionines, mutation of the periplasm-exposed Met50 did not impair function.

Collectively, our findings position RcrB as a front-line oxidative stress defense system in UPEC that neutralizes RCS before they penetrate the cytoplasm, limiting the propagation of widespread macromolecular damage and preserving metabolic and redox homeostasis. Such a membrane-centered strategy is particularly suited to counter HOCl, an extremely reactive oxidant capable of rapidly damaging proteins, lipids, and nucleic acids. In contrast, detoxification of HOSCN, the second major product of MPO, relies on the cytoplasmic reductase RclA (83). Because HOSCN is substantially less reactive and primarily targets thiols (84, 85), cytosolic detoxification may sufficient for bacterial protection. HOCl, however, reacts with a broad spectrum of cellular targets at near diffusion-limited rates, making early interception at the cell envelope essential. Therefore, RcrB represents a mechanistically distinct component of the bacterial RCS defense architecture that reduces oxidative insult at the membrane interface before it escalates into global proteotoxic and metabolic collapse. The enrichment of the RcrR regulon among UPEC isolates (47) further underscores the selective advantage conferred by this system in neutrophil-rich host environments, where localized HOCl concentrations can be substantial.

## Materials and Methods

### Bacterial strains, plasmids, oligonucleotides, and growth conditions

Bacterial strains, plasmids, and oligonucleotides used in this study are listed in **Supplementary Table S2**. Unless otherwise mentioned, bacteria were grown aerobically at 37°C in Luria Broth (LB, Millipore Sigma) and/or 3-morpholinopropane-1-sulfonic acid minimal media containing 0.2% glucose, 1.32 mM K_2_HPO_4_, and 10 μM thiamine (MOPSg) (86). Antibiotics were used when required for plasmid maintenance.

### Cloning of RcrB-sfGFP and variants

C-terminally-fused RcrB-sfGFP under the control of the native promoter of *rcrARB* was constructed by overlapping PCR. Purified PCR amplicons of the native *rcrARB* (*P*_rcrA_) promoter region, the *rcrB* gene lacking the stop codon, *sfgfp*, and the linearized vector pUA66 were ligated using the NEBuilder HiFi DNA Assembly kit (New England Biolabs), transformed into *E. coli* DH5α cells, and positive clones selected with kanamycin. RcrB-sfGFP variants with the indicated amino acid substitutions were created using the Phusion site-directed mutagenesis kit (Thermo Scientific), using the RcrB-sfGFP plasmid as template. All constructs were verified by DNA sequencing (Eurofins).

### Isolation of human neutrophils

The ethics council of the Charité Berlin (Germany) approved blood sampling, and all donors gave informed consent according to the Declaration of Helsinki. Human neutrophils were isolated by a two-step density separation as described (87). Briefly, peripheral blood was layered on an equal volume of histopaque 1119 and centrifuged at 800 x *g* for 20 min. Peripheral blood mononuclear cells and neutrophil layers were collected separately, washed with PBS containing 0.2% human serum albumin (HSA) and pelleted at 300 x *g* for 10 min. The pellet was resuspended in PBS containing 0.2% HSA, layered on a discontinuous Percoll gradient (65-85% in 2 ml layers), and centrifuged at 800 x *g* for 20 min. The neutrophil containing band was washed in PBS containing 0.2% HSA and pelleted for 10 min at 300 x *g*. Cells were counted with a CASY cell counter (OMNI Life Science).

### Live imaging of isolated neutrophils and UPEC

Isolated human neutrophils were resuspended in RPMI 1640 medium (Gibco) supplemented with 5 mM HEPES containing 0.2% HSA. Prior to seeding, cells were stained with 2.5 µM DRAQ5, a cell-permeable far-red fluorescent DNA dye (Thermo Fisher Scientific) to enable nuclear tracking. A total of 5 × 10⁵ neutrophils were seeded per well in µ-Slide 8 Well ibiTreat chambered coverslips (ibidi). For stimulation, neutrophils were challenged with human serum-opsonized Δ*rcrB* cells transformed with plasmids RcrB-sfGFP and sf-GFP, respectively, at a multiplicity of infection (MOI) of 5. Live-cell imaging was performed using an EVOS FL Auto epifluorescence microscope (Thermo Fisher Scientific), with image acquisition every 5 minutes over the course of the experiment. The DRAQ5 signal was used to monitor individual neutrophil nuclei.

### Cellular localization of RcrB by fluorescent microscopy

Overnight LB cultures were diluted into MOPSg to an OD_600_= 0.04 and grown until early exponential phase (OD_600_= ∼0.25) before harvesting. Cells were incubated with 10 µg/mL DAPI and 5 µg/mL FM4-64 for 15 min in the dark. 3 µL of stained cells were spotted on a rectangular (24 x 50 mm) coverslip, and a 1% agarose pad (prepared in PBS) was gently placed on top to trap the cells. Images were taken using the sequential channel of Leica SP8 Confocal microscope 63x oil lens in lightening features (zoom 3, pinhole 0.68) and Z-stacks at ∼200 nm intervals. Stack images were then maximally projected, and representative data are shown here. For each biological replicate, the laser intensity and gain of each channel were constant across different samples.

### Cellular localization of RcrB by subcellular fractionation and immunodetection

Overnight cultures of CFT073Δ*rcrB* cells harboring RcrB-sfGFP were diluted into 100 mL MOPSg to an OD_600_= 0.04, grown until early-log phase (OD600= ∼0.25), and harvested by centrifugation at 11,000 x g for 5 min. Next, cells were dissolved in TE buffer, frozen at −80 °C, thawed on ice, and incubated for 30 min after adding 1 mg/mL lysozyme, 1x Halt Protease Inhibitor (Fisher), and 50 U Benzonase Nuclease (Millipore Sigma), and then lysed using 0.5 mm glass beads (BioSpecs) at 1,400 rpm, 8 °C for 30 min. Lysates were spun at 1,000 x g for 1 min to remove remaining intact cells, followed by centrifugation at 100,000 x *g* for 1 hr in a Beckman Coulter Optima-max ultracentrifuge to separate membrane and soluble fractions. The pellet containing the membrane was dissolved in 100 µL of 1% n-Dodecyl-β -D-Maltopyranoside (DDM) buffer (300 mM NaCl, 50 mM Tris-HCl, pH 7.5) at 4 °C for 30 min. Protein concentrations of the membrane fraction were determined by the Bradford assay (Coomassie protein assay reagent; Thermo Scientific) with bovine serum albumin as the standard. 750 μL of protein at a concentration of 1 mg/mL was applied onto a sucrose gradient consisting of 20%, 40%, and 53% sucrose solution in 300 mM NaCl, 50 mM Tris-HCl, pH 7.5. In a 3.5 mL tube, 33% of the total volume was filled 53% sucrose, 33% of the total volume with 40% sucrose, and the remaining 33% of the total volume with 20% sucrose. Finally, the membrane protein samples were loaded on top of the sucrose layers. The gradient was created in a Beckman Coulter Optima-max ultracentrifuge at 37,000 rpm for at least 17 hrs. Samples were harvested in 11 fractions from top to bottom and boiled at 95 °C for 5 min in reducing SDS buffer. Samples were separated by SDS PAGE using 12% SDS-polyacrylamide gels, transferred to the PVDF membrane using the iblot transfer system, and immunoblotted using α-GFP primary antibody (1:2,000 dilution) (Abcam), followed by α-rabbit horseradish peroxidase-conjugated IgG secondary antibody (1:1,000 dilution) (Promega). α-OmpF and α-SecD antibodies were used as markers for outer and inner membranes, respectively.

### Global differential gene expression analysis by RNAseq and bioinformatic analysis

Total RNA was isolated from HOCl-treated and untreated CFT073 and Δ*rcrB* cells for differential gene expression analysis. In brief, overnight cultures of CFT073 and Δ*rcrB* were diluted into 30 mL MOPSg to a final OD_600_= 0.04 and cultivated at 37°C with aeration. At OD_600_ ∼0.25, the cultures were split into 125 mL flasks to a final volume of 15 mL and either left untreated or treated with 0.75 mM HOCl. Transcription was stopped by the addition of 100% ice-cold methanol after 20 min of treatment. Total RNA was extracted using the RNA extraction kit (Macherey & Nagel), residual genomic DNA removed using the TURBO DNA-free kit (Thermo Scientific), and rRNA was depleted using the Illumina Ribo Zero Kit (Illumina) for Gram-negative bacteria. RNA sequencing was performed on an Illumina HiSeq 2500 by Novogene (Sacramento, USA). Differential gene expression analysis of three biological replicates, including normalization, was performed using DESeq2 (83). Data obtained from the reads was analyzed in the bioinformatics platform Galaxy (84). Using Sailfish, reads were mapped to the UPEC strain CFT073 reference sequence (GCA_000007445.1). Then, the number of reads mapped to each gene was counted using DESeq2 (Galaxy tool). Log2 fold changes value of the differentially regulated genes for both WT and mutants were then converted to fold changes (FC) and divided as WT/ mutant; or mutant/ WT. P values were converted to log10 fold changes. Ratio >1.5 in FC was plotted, with values >2 was highlighted. The volcano plot was created using GraphPad Prism.

### Transcriptional analysis using quantitative real-time PCR (qRT-PCR)

qRT-PCR was performed to analyze differences in the relative transcript abundance between CFT073 wildtype and Δ*rcrB* cells. Overnight LB cultures of CFT073 and Δ*rcrB* were diluted into MOPSg to an OD_600_= 0.04 and grown to mid-log phase (OD_600_= ∼0.3) before the cultures were left untreated or treated with the indicated concentrations of HOCl for 20 min. Transcription was stopped by the addition of ice-cold methanol, 1 mL cells harvested, total RNA was extracted (RNA extraction kit, Macherey & Nagel), residual genomic DNA removed (TURBO DNA-free kit, Thermo-Scientific), and mRNA reverse transcribed into cDNA using the PrimeScript cDNA synthesis kit (Takara). Expression of the indicated genes was analyzed by RT-qPCR following the manufacturer instructions for Radiant Green Lo-ROX (Alkali Scientific), and expression levels normalized to the 16S rRNA-encoding housekeeping gene *rrsD*. Fold changes in gene expression were calculated using the 2^ΔΔCT^ method (88).

### Detection of protein carbonylation

CFT073 and Δ*rcrB* cells were grown aerobically in MOPSg media until the early-log phase (OD_600_= ∼0.25). Cultures were then treated with 0.8 mM HOCl for 60 min. Cells equivalent to 0.5 mL of OD_600_= 1 were harvested by centrifugation and resuspended in 100 μL TE buffer containing 0.1 mg/mL lysozyme, 25 U benzonase, and 1x HALT protease inhibitor cocktail (Thermo Scientific). Following a quick freeze-thaw cycle, cell samples were lysed using 0.5 mm glass beads in the presence of carbonylation extraction buffer (Abcam). 100 μL of cell lysate was pelleted by centrifugation and dissolved in 10 μL TE buffer. 5 μL of the cell pellet was then derivatized using Abcam protein carbonylation kit and visualized after immunoblotting following the manufacturer’s instructions.

### Quantification of IbpA-msfGFP foci formation by fluorescence microscopy

MG1655 cells with chromosomally tagged IbpA-msfGFP were used to analyze HOCl-induced protein aggregation in the absence and presence of plasmid-encoded RcrB similar to as it was described before (89, 90). MG1655 IbpA-msfGFP was transformed with plasmids RcrB-pET28a and RcrB-pBAD18b, respectively. The empty vectors pET28a and pBAD18b served as controls, respectively. Cells were grown in MOPSg until early-log phase (OD_600_= ∼0.25) before HOCl was added at the indicated concentrations. At select time points, 3 µL of cells were harvested and analyzed using the Leica SP8 Confocal microscope after staining with 10 µg/mL DAPI and 5 µg/mL FM4-64 for 15-30 min. *Z*-stacks at ∼200 nm intervals were acquired. Foci formation was blind counted manually.

### Quantification of Gam-GFP foci formation by fluorescence microscopy

HOCl-induced DNA damage was examined by quantifying Gam-GFP foci over time in an engineered *E. coli* MG1655 strain carrying a chromosomal version of *gam-gfp* similar to as it was described before (89, 90). Exponentially growing cultures of MG1655 GamGFP transformed with plasmids RcrB-pET28a and pET28a, respectively, were cultivated with 100 ng/mL doxycycline to induce the expression of GamGFP and either left untreated or treated with 1 mM HOCl. Treatment with 25 µg/mL ciprofloxacin served as a positive control. 250 µL cells were harvested and washed after 1.5 hrs of treatment, incubated 15 min in dark after addition of 10 µg/mL DAPI and 5 µg/mL FM4-64. Samples were microscopically analyzed using Leica SP8 Confocal microscope. Lightening features with zoom 3, pinhole 0.68 and *Z*-stacks at ∼200 nm intervals were taken. Stack images were then maximally projected. Gam-GFP foci formation was blind counted manually.

### Detection of lipid peroxidation

Lipid peroxidation was determined using the thiobarbituric acid-reactive substances (TBARS) assay (Cell Biolabs) as described before (91, 92). In brief, exponentially growing CFT073 and Δ*rcrB* cells were either left untreated or treated with 1 mM of HOCl for 30 min. 1 ml cells were harvested at an OD_600_= 1, washed and resuspended in PBS to a final volume of 500 μL. 100 μL samples were incubated with 100 μL of SDS and 250 μL of TBA buffer (Cell Biolabs) at 95 °C for 45 min. After cooling down, samples were centrifuged at 3,000 x g for 15 min, and 200 μL was used to measure absorbance at 532 nm. The absorbance of the sample was compared to a standard curve obtained using the malondialdehyde (MDA) provided by the vendor.

### ATP quantification

The impact of HOCl on cellular ATP level in strains CFT073 and Δ*rcrB* was examined using the BacTiter Glo cell viability kit (Promega). Exponentially growing cells were treated with 1 mM HOCl. After 30 min, cells were washed twice and resuspended in PBS. Cells and the assay reagent were mixed in 1:1 ratio in a white 96-well plate, incubated with shaking for 3 min before luminescence was determined. A freshly prepared ATP standard curve was used for quantification.

### Quantification of HOCl in the media

To test the quenching ability of RcrB and its variants, the remaining amounts of HOCl present in the media of cultures with and without RcrB variants were quantified following previously published protocols (93). Δ*rcrB* containing the indicated RcrB-pBAD18b plasmid was grown in the presence or absence of 0.2% arabinose and treated with 1.25 mM HOCl for 15 min. The supernatant of 100 µL culture was incubated with 5 mM taurine (prepared in PBS) for 5 min to trap HOCl and chloramines. 25 µL of 100 mM Na-acetate buffer (pH 7.0) supplemented with 100 µM NaI were added followed by 25 µL of Tetramethylbenzidine (TMB) developer solution. Formation of oxidized TMB was quantified at 650 nm in a Tecan 200 plate reader and normalized to a taurine chloramine standard curve.

### Growth curve-based HOCl sensitivity assays with UPEC strains

The effect of HOCl on the lag phase of the indicated UPEC strains was determined in growth curve-based assays as previously described (47, 48). Overnight LB cultures of the indicated strains were diluted 25-fold into MOPSg media, cultivated until early stationary phase, and back-diluted again into fresh MOPSg to an OD_600_= 0.02. Growth was monitored under shaking conditions at 37 °C in the absence and presence of the indicated concentrations of HOCl in a Tecan Infinite 200 plate reader. Lag phase extension (LPE) was analyzed after subtracting A_600nm_= 0.3 of the untreated culture from A_600nm_= 0.3 of the HOCl-treated sample. For K-12 strains, LPE was calculated by subtracting A_600nm_= 0.15 of the untreated culture from A_600nm_= 0.15 of the HOCl-treated.

### Growth curve-based HOCl sensitivity assay of *E. coli* strains cultivated in sterile-filtered supernatants from HOCl-treated bacteria with and without RcrB

Exponentially growing CFT073, Δ*rcrB*, as well as MG1655 and Δ*rcrB* cultures harboring RcrB-pBAD18b or the EV control pBAD18b grown in MOPSg media supplemented with or without 0.2% arabinose were pelleted and resuspended in fresh MOPSg containing 1 mM HOCl for 30 min. 0.5 mL culture was harvested and the supernatant sterile-filtered using SpinX tube filters (Costar). MG1655 or MC4100 cultures were diluted into the sterile-filtered supernatant to an OD_600_= 0.02 and growth monitored in a Tecan 200 plate reader. Supernatants collected from untreated cultures were used as control. LPE was analyzed after subtracting A_600nm_= 0.3 of the untreated culture from A_600nm_= 0.3 of the HOCl-treated sample.

### Co-cultivation assays

Overnight cultures of Δ*rcrB* and Δ*rcrB*+RcrB-pBAD18b were diluted 20-fold into MOPSg supplemented with 0.2% arabinose and grown at 37 °C under shaking conditions. Once OD_600_= 0.1 was reached, cells were either kept as mono-cultures or mixed in a ratio of 1:9 (*i.e.;* ΔrcrB:Δ*rcrB*+RcrB-pBAD18b). The cultures were either left untreated or treated with 1.25 mM HOCl for 135 min. After the indicated time intervals, cells were serially diluted in PBS. Ten µL of monocultures were plated onto LB while co-cultures were plated onto LB with and without 200 µg/mL ampicillin. Plates were incubated overnight at 37 °C for colony forming units (CFUs) counts.

### Analysis of RcrB turnover during HOCl stress

Exponentially growing Δ*rcrB* +RcrB-sfGFP cells were exposed to 1 mM HOCl for 30 min prior to the addition of the indicated concentrations of chloramphenicol. At the indicated time points, cells were 10-fold diluted in 1x sterile PBS and fluorescence intensities quantified by FACS analysis using the FITC channel (PMT 50.03) of the BD FACS-Melody instrument. At least 10,000 cells were counted for each sample.

### Computational prediction of RcrB structure

RcrB amino acid sequence was obtained from KEGG GENES search and run through AlphaFold web portal (https://alphafold.ebi.ac.uk/) to predict structure. The predicted 3D model was visualized and compared using ChimeaX (94).

### Sequence analysis

Amino acid sequences of RcrB homologs from 76 bacterial strains were extracted from NCBI BLAST. Alignments of the extracted sequences were performed using ClustalW and the aligned file was then visualized with WebLogo.

## Acknowledgments

This work was supported by the NIH/NIAID grants R15AI164585-01 and R03AI174033-02 (to J.-U. D.). S.S. was supported by Weigel grants from the Phi-Sigma Biological Sciences Honors Society, a Mockford-Thompson fellowship, and a Dissertation Completion grant. P.O.T were awarded a Weigel grant from Phi-Sigma Biological Sciences. C. J. and M. B. were supported by the Illinois State University Undergraduate Research Support Program. L.I.L. and L.R.K. acknowledge funding from the German Research Foundation (DFG) through grants LE2905/2 and KN1580/1. We further thank the ISU Confocal Microscopy Facility, which was funded by NSF grant DBI-1828136. We are thankful to Rosenberg lab (Baylor College of Medicine) for the MG1655 GamGFP strain, Aertsen lab (KU Leuven, Belgium) for the MG1655 IbpA-msfGFP strain, Dr. Christopher Lennon (Murray State University) for the sfGFP plasmid, Shilhavy lab (Princeton University) for the anti-OmpF antibody, and Collinson lab (University of Bristol, UK) for the anti-SecE antibody. We are also grateful to Dr. Ursula Jakob (University of Michigan) for her willingness to provide feedback on this study. Finally, we thank present and past members of Dahl lab for their valuable feedback on the study.

