## Supplementary Figures & Tables for "A newly identified detoxification system protects uropathogenic *Escherichia coli* from reactive chlorine species"

### Supplementary Results

**RcrB Does Not Alter Membrane Integrity or Polarization.** Because RcrB localizes uniformly to the inner membrane (**Fig. 1D**) and loss of RcrB leads to elevated lipid peroxidation (**Fig. 3F**), we tested whether its protective effect reflects other membrane characteristics. Outer and inner membrane integrity were assessed using N-phenyl-1-naphthylamine (NPN) and propidium iodide (PI), respectively, the latter of which enters cells only when the inner membrane is compromised (1). Exposure to sublethal HOCl modestly increased PI fluorescence in both strains, indicating at least partial HOCl-induced membrane perturbations (*SI Appendix, Fig. S3A*). In contrast and unlike polymyxin B, which served as a positive control for outer membrane disruption, NPN fluorescence remained unchanged following HOCl treatment (*SI Appendix, Fig. S3B*). Importantly, no significant differences in PI or NPN uptake were observed between WT and Δ*rcrB* cells, indicating that RcrB has no effect on basal or HOCl-induced membrane permeability. We next examined membrane polarization using the $DiOC_2$, a membrane potential-sensitive dye (2). HOCl exposure caused a comparable concentration-dependent increase in $DiOC_2$ fluorescence in both WT and Δ*rcrB* cells, indicating RcrB-independent membrane hyperpolarization upon HOCl stress (*SI Appendix, Fig. S3C*). Thus, RcrB does not modulate HOCl-induced changes in membrane physiology.

### Supplementary Material and Methods

**RcrB-sfGFP expression by flow cytometry.** Overnight LB cultures were diluted into MOPSg to an $OD_{600}$= 0.04 and grown until early exponential phase ($OD_{600}$= ~0.25) before 0.75 mM HOCl was added. At the indicated time points, cells were 10-fold diluted in 1x sterile PBS and fluorescence intensities quantified using the FITC channel (PMT 50.03) of the BD FACS-Melody instrument. At least 10,000 cells were counted for each sample.

**Quantification of IbpA-msfGFP expression by flow cytometry.** MG1655 cells with chromosomally tagged IbpA-msfGFP was used to analyze HOCl-induced protein damage in the absence and presence of plasmid-encoded RcrB expression. MG1655 ibpA-msfGFP was transformed with plasmid RcrB-pET28a. The empty vector pET28a served as control. Cells were grown in MOPSg until early-log phase ($OD_{600}$= ~0.25) before HOCl was added at the indicated sublethal concentrations. At the indicated time points, cells were 10-fold diluted in 1x sterile PBS and fluorescence intensities quantified using the FITC channel (PMT 50.03) of the BD FACS-Melody instrument. At least 10,000 cells were counted for each sample.

**Luciferase reporter assay.** Overnight MOPSg culture of CFT073 and Δ*rcrB* strains containing *PsulA-luxCDABE* were diluted into fresh MOPSg to an $OD_{600}$ of 0.25. 190 µL of bacterial culture were added per well in a 96-well plate and treated with the indicated concentrations of HOCl. Luminescence and $OD_{600}$ were measured for 8 hrs in 20 min intervals using a Tecan plate reader.

**Quantification of $NAD^+$, NADH, $NADP^+$ and NADPH.** UPEC strains CFT073 and Δ*rcrB* were grown in MOPSg until mid-exponential phase and treated with 1 mM HOCl. After 30 min, 2 mL cells equivalent of $OD_{600}$= 0.5 were harvested, washed, and resuspended in PBS. $NAD^+$/NADH and $NADP^+$/NADPH levels were measured following the manufacturer's instructions for the respective Glo assay kits (Promega). For detection of $NAD^+$ and $NADP^+$, cells were lysed with 0.2 M HCl at 60ºC for 15 min and neutralized to pH 7.0 with NaOH. For detection of NADH and NADPH, cells were lysed with 0.2 M NaOH at 60ºC for 15 min and neutralized to pH 7.0 with HCl. Cell lysates were incubated with the respective assay master mix in 1:1 ratio. $NAD^+$, NADH, $NADP^+$, and NADPH concentrations were determined by luminescence measurements and quantified through a standard curve with the corresponding nicotinamide.

**Quantification of inner membrane and outer membrane damage.** As described before (86), propidium iodide (PI) and N-phenyl-1-naphthylamine (NPN) fluorescence was used to quantify damage of the inner and outer membrane, respectively. Exponentially growing CFT073 and Δ*rcrB* cells were incubated with the indicated concentrations of HOCl for 30 min, washed and resuspended in PBS containing 5 µM PI and 10 µM NPN, respectively. After 15 min incubation in the dark, fluorescence was measured in the Tecan 200 microplate reader at exc./em. wavelengths of 535/613 nm for PI and 350/420 nm for NPN, respectively. Changes in NPN fluorescence were calculated using the equation in (87).

**Analysis of the membrane potential.** Changes in membrane potential were quantified using the 3,3'-Diethyloxacarbocyanine Iodide (DiOC2) fluorescent probe. Exponentially growing CFT073 and Δ*rcrB* cells were either left untreated or treated with the indicated HOCl concentrations, harvested after 30 min and resuspended in 250 µL PBS containing 10 mM EDTA at an $OD_{600}$= 1. After 10 min of incubation, cells were pelleted and resuspended in 250 µL assay suspension buffer supplemented with 30 µM DiOC2 (Thermo Scientific). 5 µM of the proton uncoupler carbonyl cyanide m-chlorophenylhydrazone (CCCP) was used as a control to depolarize the membrane. After 15 min incubation in the dark, cells were washed with PBS and fluorescence intensities measured in the PerCP-Cy5.5 channel of the BD FACS Melody flow cytometer. Mean fluorescent intensities were quantified using FCSalyzer0.9.18.

### Supplementary Figures and Tables

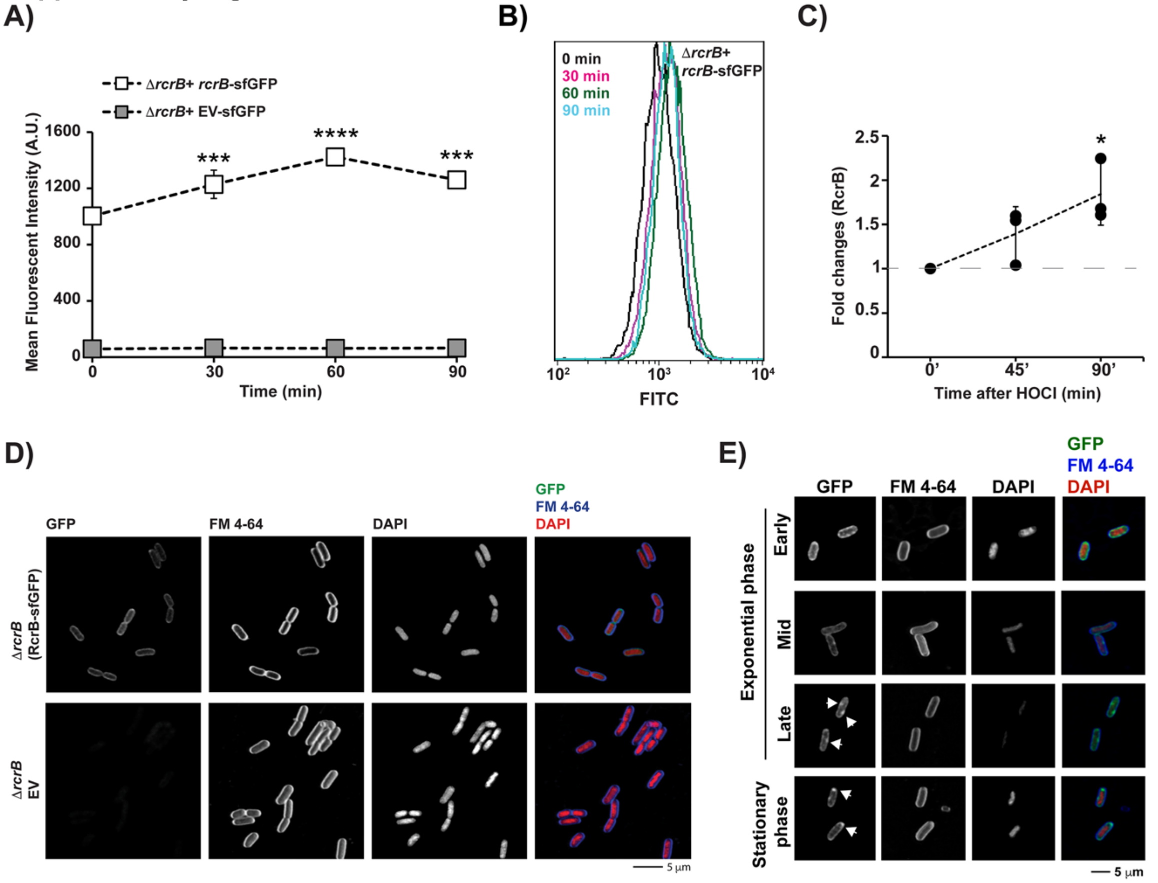


**Supplementary Fig S1: Expression of the membrane protein RcrB is induced upon HOCl stress. (A,B)** Exponentially growing Δ*rcrB* cells with and without RcrB-sfGFP expression were treated with 0.75 mM HOCl. GFP fluorescence intensities were quantified using flow cytometry. **(A)** shows the mean fluorescence intensities of four independent experiments (n= 4, mean±S.D.), and **(B)** a representative micrograph of the fluorescent intensity values of Δ*rcrB* + RcrB-sfGFP cells after treatment with 0.75 mM HOCl. **(C)** Quantification of RcrB-sfGFP protein level using ImageJ analysis revealed a ~2-fold increase after 90 min of HOCl exposure (n=3, mean±S.D.). **(D)** Representative confocal micrograph of four biological replicates shows the localization of RcrB-sfGFP (green) in the membrane of exponentially growing CFT073Δ*rcrB*. DAPI (red) and FM4-64 (blue) were used to stain DNA and membrane, respectively. The right panel shows an overlay image of GFP, FM 4-64, and DAPI fluorescence. **(E)** Localization and expression pattern of RcrB-sfGFP in UPEC CFT073Δ*rcrB* cells at different growth stages. One representative image of three independent experiments is shown. One-way ANOVA (Dunnett's post-test) between different cell samples (GraphPad Prism) *p<0.05; ***0.001>p>0.0001; ****p<0.0001.

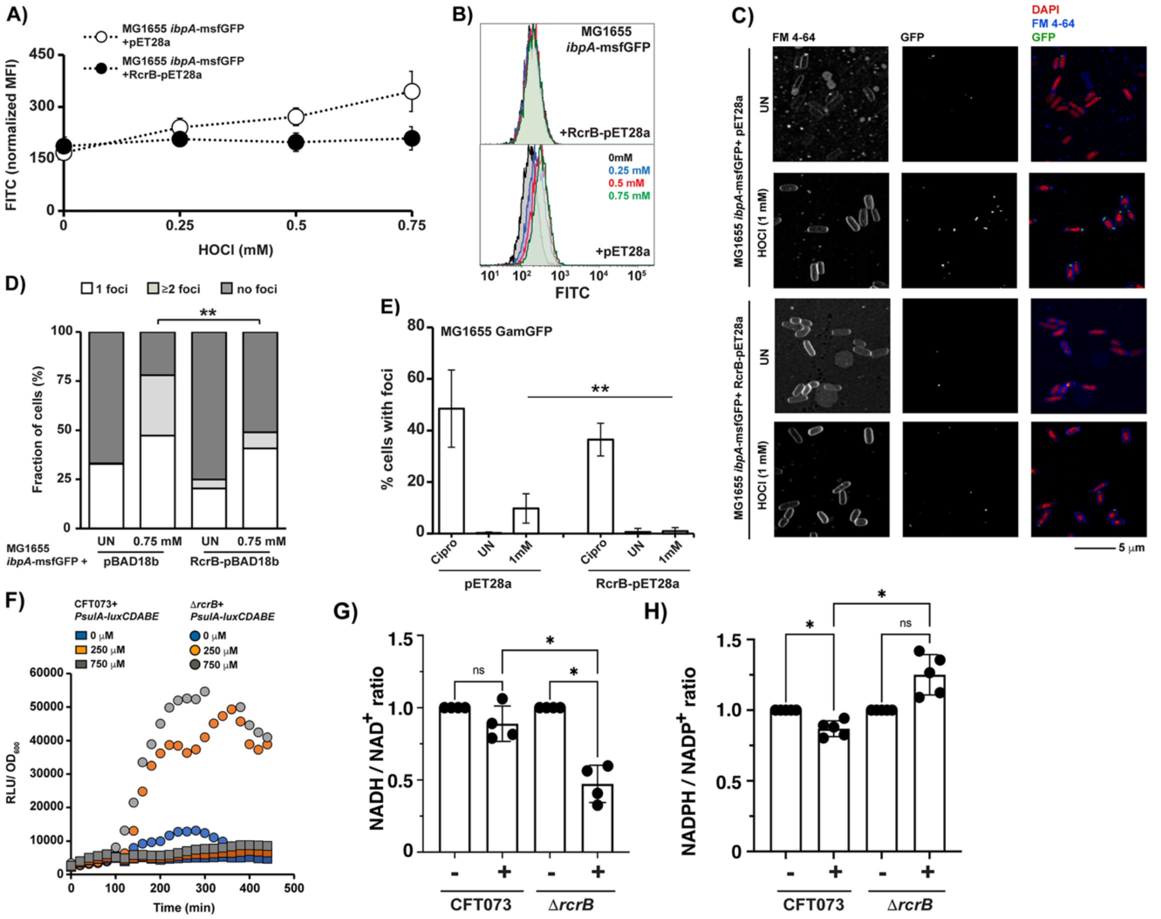


**Supplementary Fig S2: Expression of RcrB protects cells from HOCl-induced damage. (A, B)** Exponentially growing MG1655 encoding a chromosomally fused IbpA-msfGFP fusion were transformed with the plasmid RcrB-pET28a and the EV control, respectively. Cells were treated with the indicated HOCl concentrations for 90 min before IbpA-msfGFP fluorescence was determined by flow cytometry. **(A)** shows the mean fluorescence intensities of four independent experiments (n=4, mean±S.D.), and **(B)** representative micrographs from the FACS. **(C)** MG1655 *ibpA*-msfGFP cells transformed with either RcrB-pET28a or the EV control were grown in MOPSg to early exponential phase and incubated with HOCl for 30 min. IbpA-msfGFP foci formation (green) was then visualized by confocal microscopy using Leica SP8. DAPI (red) and FM4-64 (blue) are used to stain DNA and outer membrane, respectively. One representative image of four biological replicates is shown. **(D)** MG1655 *ibpA*-msfGFP cells transformed with either RcrB-pBAD18b or the pBAD18b EV control were grown in MOPSg to early exponential phase and incubated with 1 mM HOCl for 30 min. IbpA-msfGFP foci formation was then visualized by confocal microscopy. ImbpA-msfGFP foci were counted blindly (n=4, mean±S.D.). **(E)** Exponentially growing cells of the MG1655 Gam-GFP reporter strain carrying either RcrB-pET28a or the EV control were treated with 1 mM HOCl and 25 µg/mL ciprofloxacin (Cipro) for 2 hrs. Gam-GFP foci formation was visualized under the microscope indicative of double-strand DNA breaks. Fluorescent foci were counted blindly. (n=5, mean±S.D.). **(F)** Representative luminescence plot of sulA promoter activity of four independent experiments. Exponentially growing CFT073 and Δ*rcrB* cells with sulA-promoter luciferase fusion ($P_{sulA}$-*luxCDABE*) were incubated with the indicated concentrations of HOCl. Luminescence intensities were measured over 480 min using a Tecan 200 plate reader. **(G&H)** Reduced and oxidized nicotinamide adenine nucleotide levels were analyzed in strains CFT073 and Δ*rcrB* before and after treatment with 1 mM of HOCl for 30 min.

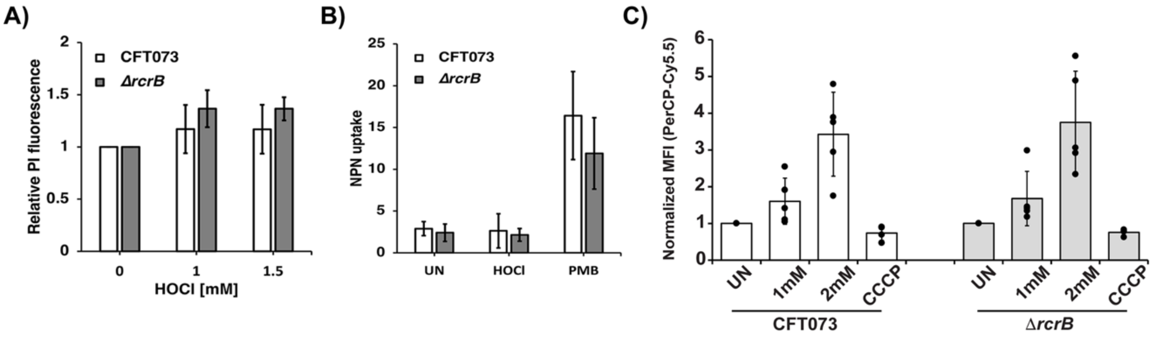


**Supplementary Fig S3: Expression of RcrB during HOCl stress does not affect the cell envelope integrity. (A,B)** CFT073 and Δ*rcrB* were grown to the mid-log phase in MOPSg media and either left untreated or treated with the indicated concentrations of HOCl for 30 min. Cells were harvested, washed in PBS, and stained with **(A)** 0.5 µM PI and **(B)** 10 µM NPN dye. Fluorescence intensities were determined at excitation/emission wavelengths of **(A)** 535/617 nm and **(B)** 350/420 nm, respectively. (n=4-5, mean±S.D.) **(C)** Changes in membrane potential were quantified using the DiOC2 fluorescent probe after incubating CFT073 and Δ*rcrB* for 30 min with the indicated HOCl concentrations. 5 µM CCCP was used as a depolarization control (n=5, mean±S.D.).

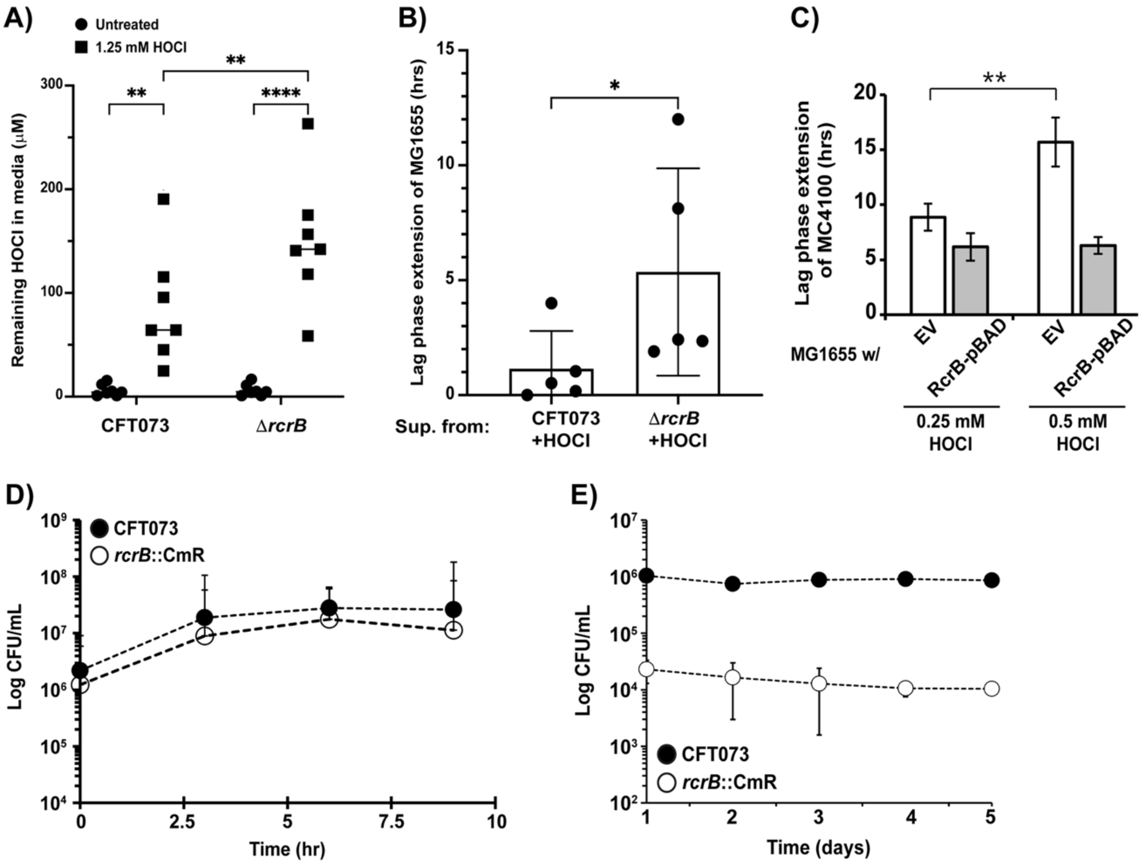


**Supplementary Fig S4: RcrB plays a role for HOCl detoxification. (A)** Detection of HOCl in the supernatant of exponentially growing CFT073 and Δ*rcrB* cultures stressed with 1.25 mM HOCl for 15 min was performed using the taurine chloramine (TMB) assay (n=7, mean±S.D.). **(B)** Lag phase extension of *E. coli* K-12 strain MG1655 was determined during incubation with spent media from CFT073 and Δ*rcrB* cells that had been exposed to 1 mM HOCl for 30 min. (n=5 ± S.D.). **(C)** Lag phase extension of *E. coli* K-12 strain MC4100 was determined during incubation with spent media from MG1655 cells containing the RcrB-pBAD18b expression plasmid and the pBAD18b EV control, respectively, which had been exposed to the indicated HOCl concentrations for 30 min. (n=4 ± S.D.). **(D,E)** CFT073 and CFT073Δ*rcrB*::$Cm^R$ were co-incubated **(A)** in a 1:1 ratio for 9 hrs, or **(B)** in a 9:1 ratio for five days to examine the competitive fitness of Δ*rcrB* under non-stress conditions. (n=4, mean±S.D.).

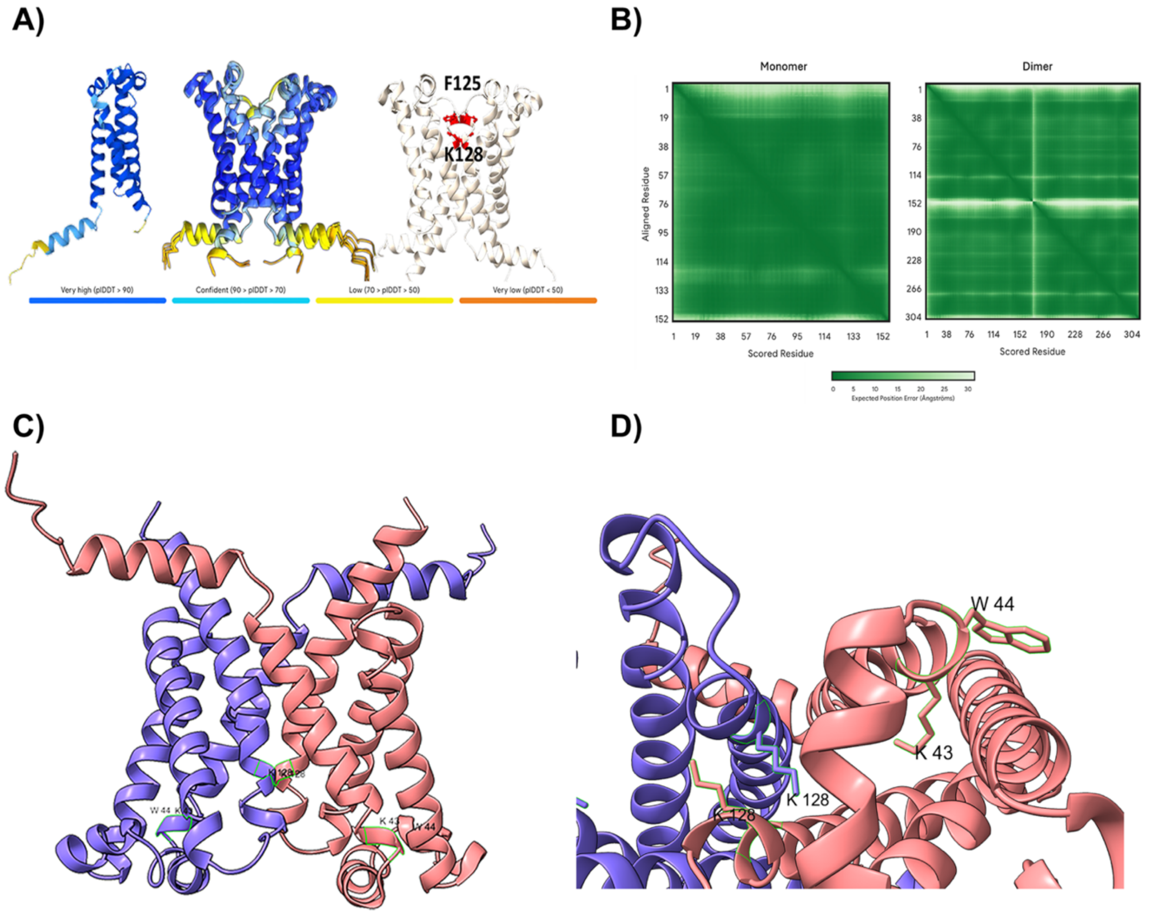


**Supplementary Fig S5: Predicted AlphaFold structure of RcrB. (A)** The predicted dimer interface is formed by hydrophobic residues within the transmembrane helix, with Lys128 and Phe125 stabilizing the periplasm-facing region of the interface. **(B,C)** Predicted AlphaFold structure of RcrB dimers. **(D)** Bottom view of the predicted AlphaFold structure of RcrB dimers.

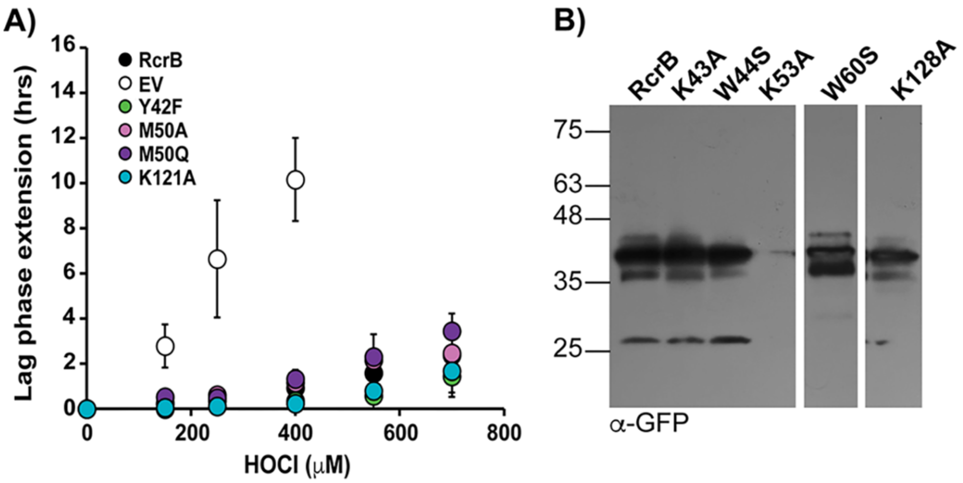


**Supplementary Fig S6: Characterization of RcrB variants with substitutions in redox-sensitive amino acids. (A)** Complementation analyses of the Δ*rcrB* strain expressing RcrB-sfGFP variants with the indicated amino acid substitutions were performed in MOPSg media in the presence of the indicated HOCl concentrations. EV= empty vector. HOCl-mediated LPE was calculated for each strain (*see Materials and Methods*) (n=3-5, mean±S.D.). **(B)** Exponentially growing Δ*rcrB* + RcrB-sfGFP cells were lysed and the membrane fraction was isolated by sucrose-gradient ultracentrifugation. RcrB-sfGFP was detected by western blot analysis using α-GFP antibody. One representative image of three independent experiments is shown.

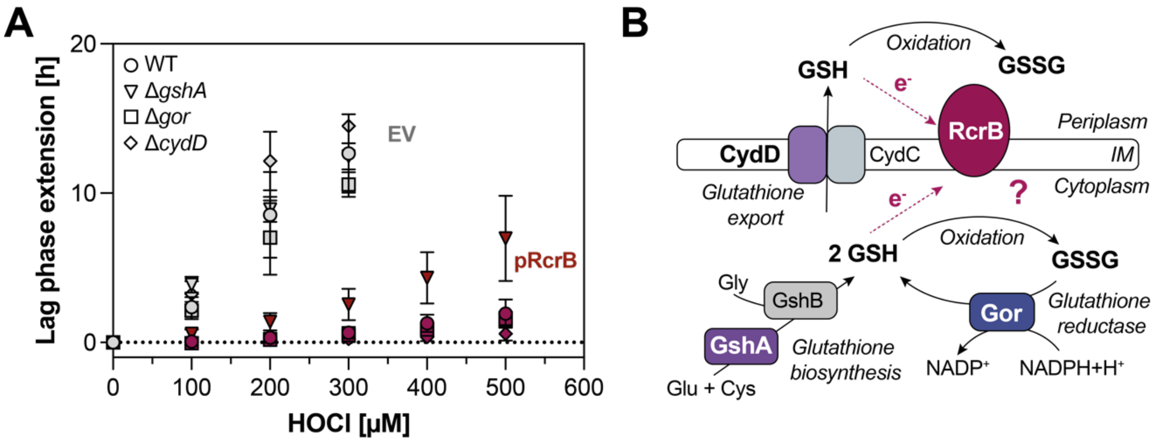


**Supplementary Fig S7: Expression of RcrB requires a functional *gshA* gene for full complementation of ∆*rcrB*. (A)** Growth analyses of the *E. coli* BW25113 and KEIO collection mutants deficient in *gshA, gor,* and *cydD* (3). Cells that either express RcrB from the pBAD18 expression plasmid (pRcrB) or contain the EV control (EV) were cultivated in MOPSg media in the presence of the indicated HOCl concentrations. HOCl-mediated LPE was calculated for each strain (*see Materials and Methods*) (n = ≥3, mean± S.D.). grey bars/symbols: pBAD18b; red bars/symbols: RcrB-pBAD18b. **(B)** Proposed model for the reduction of oxidized/chlorinated RcrB.

**Supplementary Table S1. Differentially expressed genes in HOCl-treated CFT073 and ∆*rcrB***

**Supplementary Table S2. Bacterial strains, plasmids, and oligonucleotides used in this study.**
